# Rest-fMRI Based Comparison Study between Autism Spectrum Disorder and Typically Control Using Graph Frequency Bands

**DOI:** 10.1101/2021.01.29.428745

**Authors:** Alireza Talesh Jafadideh, Babak Mohammadzadeh Asl

**Affiliations:** Department of Biomedical Engineering, Tarbiat Modares University, Tehran, Iran

**Keywords:** Autism spectrum disorder, Typical control, Diffusion tensor imaging, Graph signal processing, Resting-state fMRI

## Abstract

Graph signal processing is a subset of signal processing enabling the analysis of functional magnetic resonance imaging (fMRI) data in brain topological domain. One of the most important and highly interested tool of GSP is graph Fourier transform (GFT) by which brain signals can be analyzed in different graph frequency bands. In this paper, the resting-state fMRI (rfMRI) data is analyzed using GFT tool in order to discover new knowledge about the autism spectrum disorder (ASD) and find features discriminating between ASD and typically control (TC) subjects. For ASD group, the signal concentration in both low and high frequency bands is decreased by increasing the age in most of the brain well-known networks. The ASD in comparison to TC shows less intention for changing the signal concentration level when the level is very low or very high. In graph low frequency band, increasing the age is along with increasing the segregation and integration of brain ROIs respectively for ASD and TC. Also, in this band, the brain ROIs integration of ASD is more than TC. By increasing the age, the auditory network of ASD subjects shows increasing segregation and integration in graph low and high frequency bands, respectively. The reduced segregation of default mode network in ASD is happened in graph middle and higher frequency bands. The functional connectivity analysis between low and high frequency signals shows that some of the high frequency ROIs have connections with all low frequency ROIs so that the most of these connections are dramatically and significantly different between ASD and TC. By analyzing the local vertex frequency spectrum (LVFS) of ASD and TC at different states, it is seen these groups show contradictory behaviors with respect to each other in brain default mode network in two states. The results of different scenarios at different graph frequency bands demonstrate that using functional and structural data together can provide powerful tool for recognizing the ASD or even other brain disorders.

## 1. Introduction

Autism spectrum disorder (ASD) is a heterogeneous neurodevelopmental disorder mainly characterized by deficits in social communication and interaction, restricted interests, and repetitive behaviors or activities (American Psychiatric Association, 2013). ASD imposes severe adverse effects and economical costs on subjects, their families, medical parts, and society (Buescher et al., 2014). Unfortunately, the ASD prevalence is rapidly growing and its etiology is heterogeneous (Elsabbagh et al., 2012). Because of this heterogeneity and speed of prevalence, a lot of researches have been conducted for finding biomarkers and features in order to make possible the early and reliable diagnosis (Goldani et al., 2014; Woo and Wager, 2015; Drysdale et al., 2016; Li et al., 2017). It is accepted that this disorder makes changes in both structure and function of brain (Ha et al., 2015). As a result, investigating the structural and functional changes of ASD individuals’ brain using neuroimaging tools and methods can provide reliable biomarkers and quantifiable features for ASD diagnosis (Grecucci et al., 2017).

One of the most popular and accepted approaches for structural and functional analysis of brain is brain connectivity in which the interactions between different regions of interest ROIs are studied (Uddin et al., 2013; Ha et al., 2015; Mash et al., 2018). The most widely used modality for functional connectivity (FC) analysis is functional magneto resonance imaging (fMRI) by which the indirect measure of brain activity is given by measuring the blood oxygen level-dependent (BOLD) signal (Filler, 2009). The resting-state fMRI (rfMRI) technique has been mostly employed for FC because of its fast and task-free nature (Biswal, 2012; Hull et al., 2017). Two common approaches exist for studying the FC: the traditional static FC (SFC), the recently developed dynamic FC (DFC). The earlier one uses all time points of ROIs’ signals for computing FC value. The SFC researches have resulted in both stronger and weaker connectivity for ASD subjects in comparison to the typical control (TC) ones (Cheng et al., 2015; Iidaka, 2015; Hull et al., 2016). This conflict may be a result of overlooking the time varying nature of fMRI data (Mash et al., 2019). To overcome this problem, the latter one usually employs the sliding window (SW) technique and computes sequence of FC matrices using a subset of total time points and finally these matrices are entered into clustering process for finding the “states” (Preti et al., 2017; Rashid et al., 2018; Aggarwal and Gupta, 2019; Mash et al., 2019). It has been stated that DFC may provide answers for SFC conflict results by residing weaker and stronger connectivity of ASD in different states (Mash et al., 2019). In (Mash et al., 2019), the authors found that reduced default mode network segregation occurs by involving somatomotor network in one state and fronto-parietal and executive networks in another state. In (Rashid et al., 2018), it is found that dwelling time has positive relation to the autistic traits severity in a “globally disconnected” state.

Another type of brain connectivity is structural connectivity in which the brain ROIs are connected through a bundle of fibers. The fiber tracts are usually rebuilt using diffusion tensor imaging (DTI) tractography, and then are used for determining the strength of between ROIs connection and consequently structural connectivity matrix estimation (Mukherjee and McKinstry, 2006; Jou et al., 2011; Mash et al., 2018). Using DTI data, increased mean diffusivity (MD) has been reported in the whole frontal lobe (Sundaram et al., 2008), temporal portion of the left superior longitudinal fasciculus (Nagae et al., 2012), and thalamo-cortical connections (Nair et al., 2013) of ASD children.

In one point of view, the rfMRI data is time series signals of ROIs changing across temporal dimension. Thus, only the processing tools of this dimension might be used for analysis the behavior of this data. However, in another point of view, it can be said that the rfMRI data is resided on the brain topology in which the ROIs are connected with a structural connectivity strength. As a result, the rfMRI data can be analyzed in topological domain. In contrast to the temporal domain in which the samples are in a regular environment, the topological domain is irregular meaning the samples are not positioned on a straight line, the distances between all two adjacent connected samples can be different from each other, and each sample can have more than two adjacent connected samples. Therefore, the temporal analyzing tools such as filtering and concepts such as Fourier frequency cannot be directly employed in the topological domain. Fortunately, the graph signal processing (GSP) is an emerging field attempting to develop the regular domain tools for irregular one (Shuman et al., 2013; Ortega et al., 2018). The GSP uses two components for combinatorial usage of the brain structural and functional data: graph and signal. The graph *𝒢* consists of vertices *𝒱* and edges *ε*. In this study, the vertices are ROIs and *ε ⊂ 𝒱* × *𝒱* represents the structural connectivity between ROIs. The signal vector of GSP is the rfMRI data residing on vertices at each time repetition (TR). Therefore, the GSP technique enables the combined use of structural and functional data.

Recently, the GSP application in neuroimaging field has been investigated (Huang et al., 2016; Ménoret et al., 2017; Huang et al., 2018; Medaglia et al., 2018; Wang et al., 2018). In (Huang et al., 2018), the associate between attention switching cost and brain signal concentration was examined in graph low and high frequencies. The results showed a positive significant relation for high frequency signals, meaning that the hardest switching task for subjects were along with the high signal variance with respect to the underlying brain topology. In (Huang et al., 2016), the GSP was exploited for studying the brain behavior during performing learning tasks. It is found that brain signals in different graph frequencies show different levels of adaptability during learning. In other works, the GSP tools have been used for feature extraction in order to improve the classification performance. For example, in (Wang et al., 2018), features from graph frequencies and features from the local and global measures of functional connectivity matrix were combined for classifying different age groups. The results showed an accuracy of 86.64% for 389 subjects.. In all mentioned papers, the most important tool of GSP, i.e., graph Fourier transform (GFT) was used for analyzing the fMRI. In (Huang et al., 2018), (Huang et al., 2016; Medaglia et al., 2018; Wang et al., 2018), and (Ménoret et al., 2017), respectively, the structural, functional, and both structural and functional connectivity matrices were employed as *𝒢*. In (Ménoret et al., 2017), the Gaussian kernel of the distance between ROIs was used for computation of connectivity strength.

Recently, the GFT has been utilized in two works to classify ASD and TC (Itani and Thanou, 2019; Brahim and Farrugia, 2020) using Autism Brain Imaging Data Exchange (ABIDE) I preprocessed database (Di Martino et al., 2014; Preprocessed Connectomes Project, 2014). In (Brahim and Farrugia, 2020), the structural graph averaged across 56 TC subjects from the Human Connectome Project (HCP) was used as graph of GSP for all understudy ABIDE I subjects. This structural graph was obtained using tractography of the brain white matter. Some statistical metrics including standard deviation (SD), mean, variance, and Kurtosis were calculated for each ROI time series and used as graph signals. The mentioned statistical metrics of ROIs, their GFT, and also the eigenvector centrality, vertex strength and clustering coefficient of functional connectivity matrix were considered as features. Using the GFT of SD as feature offered the best classification performance in comparison to the other features and also other state of the art methods. In (Itani and Thanou, 2019), the structural graph was defined based on the distance between ROIs so that the strength between two ROIs was equal to inverse of their distance from each other. For each ROI, only the first two nearest ROIs were kept. The authors proposed framework used both GFT and spatial filtering method (SFM) for finding a subspace in which the discriminative features could be extracted for ASD and TC groups. In SFM, the time-series of two class instances were projected in a Fukunaga-Koontz Transform (FKT) space so that they became distinguishable. The decision tree classification results showed superiority of their proposed framework than the other state of the art methods.

In this paper, different graph frequency bands with variant techniques are investigated in order to find discriminative characteristics between ASD and TC groups. The San Diego State University (SDSU) dataset of ABIDE II database which has both DTI images and rfMRI data is used (Di Martino et al., 2017). The DTI images of each subject are employed for subject-specific structural connectivity matrix computation. In this paper, different scenarios are implemented and their measures relations to age and also their measures differences between ASD and TC are examined. The main scenarios of this paper are as follows:

1. Examining the signal concentrations through mean concentration coefficient (MCC) (Huang et al., 2018) for ach brain network in graph low and high frequencies
2. Examining the quartile CC (QCC) for total brain in graph low and high frequencies
3. Examining the SFC behavior at ROI and network levels in graph low and high frequencies
4. Examining the cross frequency FC between graph low and high frequency signals
5. Examining the local vertex frequency spectrum (LVFS) (Stankovic et al., 2019) in different states in order to analyses group differences in graph frequency domain

The rest of the paper is organized as follows: dataset description and preprocessing are given in section 2. This section is continued by explaining the GFT and graph filtering and then describing the scenarios. The final subsection of this section is devoted to statistical evaluations. The results are provided in section 3. Discussion about the results and conclusions are respectively offered in sections 4 and 5.

## 2. Materials and Methods

### 2.1. Participants and data acquisition

In this study, the SDSU dataset of ABIDE II database is exploited. This dataset includes 58 subjects (33 ASD, 25 TC) ages 7.4-18 years. In this study, the data of one TC subject (the ID number “28852” of ABIDE II) is rejected due to motion artifact problem (see section 2.2.1). Thus, the 57 subjects are studied. Two groups are matched on age, handedness, sex, and head motion while show significant difference in social responsiveness scale (SRS). The SRS of one ASD subject is missing. The sample characteristics are listed in Table 1. The root-mean-squared displacement (RMSD) is explained in the section 2.2.1.

**Table 1.** Sample characteristics.

|  | ASD (33) | TC (24) | Statistic | *p* |
| --- | --- | --- | --- | --- |
| Age (year) | 12.89±3.28<br>[7.4-18] | 13.39± 3.02<br>[8.1-17.7] | *t*(55)=-0.59 | 0.56 |
| Sex | 26 Male | 22 Male | $\chi^2(1)=1.73$ | 0.19 |
| Handedness | 27 Right | 20 Right | $\chi^2(1)=0.02$ | 0.88 |
| SRS total | 83.37± 9<br>[62-94] | 42.62± 5.21<br>[35-52] | *t*(54)=19.81 | <0.001 |
| RMSD (mm) | 0.04± 0.03<br>[0.01-0.12] | 0.03± 0.03<br>[0.01-0.11] | *t*(55)=0.39 | 0.7 |

Data collection was performed using a GE 3T MR750 scanner with an eight-channel head coil. High-resolution structural images were acquired with a standard SPGR T1-weighted sequence (TR/TE = 8.136/3.172 ms, flip angle =8 degree, field of view (FOV) = 256×256 mm, 1 mm^3^ resolution, 172 slices). The rfMRI datasets were collected using a standard gradient echo-planar imaging (EPI) sequence with TR/TE = 2000/30 ms, flip angle = 90 degree, FOV = 220×220 mm, matrix size = 64×64, pixel spacing size = 3.4375×3.4375 mm, slice thickness = 3.4 mm, slice gap =0 mm, axial slices = 42, and total volumes = 180. DTI protocol was based on a 2D spin-echo EPI sequence. DTI images, consisting of 60 weighted diffusion scans (b = 1000 sec/mm^2^) and one unweighted diffusion scan (b = 0 sec/mm^2^), were recorded with the following parameters: TR/TE = 8500/78 ms, FOV = 128×128 mm, slice in-place resolution 0.9375×0.9375/1.875×1.875 mm^2^, slice thickness = 2 mm, 68 axial slices.

### 2.2. Data preprocessing

#### 2.2.1. rfMRI preprocessing and time series extraction

The rfMRI data was preprocessed using SPM^1^ and AFNI^2^ software packages and also personal codes. In this study, the first five volumes were discarded to account for T1-equilibration effects. For each subject, the typical preprocessing steps including slice-timing correction using the middle slice as the reference time frame, motion correction, despiking using AFNI’s 3dDespike, coregistration of T1 and functional images, warping images to the standard Montreal Neurological Institute (MNI) space, and spatially smoothing using a Gaussian kernel with a 5 mm full width at half maximum (FWHM = 5 mm) were implemented.

Some post-preprocessing steps were performed in order to reduce the noise. Six motion parameters and their first temporal derivatives and one principle component of cerebrospinal fluid (CSF) and white matter (WM) signals were considered as 14 confounds and regressed out from all voxels time series. For finding the CSF/WM confounds, the CSF/WM mask available in WFU PickAtlas software^3^ was employed for extracting time series of its voxels. Then, the principle components of CSF/CM were attained by principle component analysis (PCA) method. For each of the CSF and WM, one dominant component explained more than 99% time series variance and so they were considered as confounds (Caballero-Gaudes and Reynolds, 2017). The preprocessing was continued by detrending the linear, quadratic, and cubic trends, and then applying band-pass fifth-order Butterworth filter (0.01-0.1 Hz) on the time series.

Some of the volumes were possible to be corrupted due to motion artifacts. For detecting them, the differences between two consecutive time points motion parameters were computed (totally 6 difference values for six motion parameters). Then, the square root of the sum of squares of computed differences was regarded as framewise displacement (FD). For transforming radians of rotational parameters to millimeter, the head was regarded as a sphere of radius 50 mm (Power et al., 2012). The time points with FD >0.5 mm and also the two subsequent time points were censored (Mash et al., 2019). As with (Mash et al., 2019), time series fragments with <10 consecutive time points remaining after censoring were also excluded. After censoring corrupted time points, if the number of remaining time points of subject was less than 75% of the number of the initial time points (175), that subject was rejected for using in this study. The TC subject with ID = 28852 was rejected. The average of eliminated time points for ASD and TC were 0.95 and 0.45, respectively. 25 ASD and 22 TC subjects had no censoring time points meaning no significant difference between two groups (χ2(1) = 1.73, *p =* 0.19). The maximum censored time points for one subject of ASD and TC were 19 and 16 which respectively were 34.54% and 84.21% of the total number of censored points of these groups. Two groups didn’t show significant difference in term of the number of eliminated time points (t(55) = 0.82, *p =* 0.41). It is more convenient to have equal number of time points for all understudy subjects. Hence, by considering that the maximum number of censored points and the initial time points respectively were 19 and 175, the first 156 time points for each subject were selected for further analysis (Goldani et al., 2014).

Based on the MNI coordinates of regions of seven networks including default mode network (DMN), cognitive control network (CCN), auditory network (AN), somatomotor network (SMN), visual network (VN), cerebral network (CN), subcortical network (SCN) reported in (Rashid et al., 2018; Allen et al., 2014), and based on the MNI coordinates of 116 ROIs of automated anatomical labeling (AAL) atlas, 98 out of 116 ROIs of AAL atlas were chosen and assigned to 7 aforementioned networks (see Supplementary Table S1). For ASD subject with ID = 28869, the “Cerebelum_7b_R” didn’t show any structural connectivity with other ROIs (see section 2.2.2). Thus, this ROI was rejected from our analysis. By averaging the time series of ROI voxels, one signal was obtained for each ROI. Therefore, the final rfMRI data is 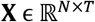 where *N = 97* and *T = 156* are the numbers of ROIs and time samples, respectively.

#### 2.2.2. DTI preprocessing and structural connectivity matrix

In this study, the ExploreDTI software was used for DTI data preprocessing and structural connectivity matrix computation. After performing some initial steps for data preparation and quality assessment in ExploreDTI software, diffusion data were corrected for subjects’ motion and Eddy currents. Then, deterministic fiber tracking was performed for tractography purpose. Using the tractography result and AAL atlas, the structural connectivity matrix was computed for each subject. In this matrix, the connectivity strength between two regions was equal to the number of tracts existing between them. In the next step, each row of this matrix was divided to sum of its elements. By doing so, the value of entry (*i,j*) expressed the connectivity probability from region *i* to region *j*. The probability from *i* to j was not equal to that from *j* to *i*. Thus, the connectivity matrix which is also called the adjacency matrix **A** was not symmetric. As the final step, the symmetric adjacency matrix **A’** was obtained as **A’ =** (**A + A^T^**)/2 where.^T^ denotes the transpose operator.

### 2.3. Graph frequencies and filtering

One of the most important GSP tool is GFT by which the graph frequency domain and graph frequencies are defined. Let us define the combinatorial Laplacian matrix 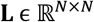 which is the fundamental matrix for implementing GFT (Shuman et al., 2013) as follows:

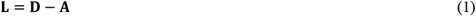

where **D** is a diagonal matrix including the degrees of each vertex and it’s *k*^th^ diagonal element is 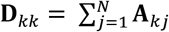. The Laplacian matrix is real, symmetric, and positive semi-definite (Von Luxburg, 2007). Thus, it can be eigen-decomposed as follows:

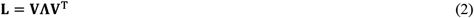

where Λ is diagonal eigenvalues matrix with *k*^th^ diagonal element of *λ_k_* and **V** is eigenvector matrix so that **V**^Γ^**V** = **I**_*N*_ and **I**_*N*_ represents the N-dimensional identity matrix. The sorted eigenvalues of graph Laplacian are 0 = *λ*_0_ < *λ*_1_ ≤ *λ*_2_ …. ≤ *λ*_*N*-1_ = *λ_Max_* and their corresponding eigenvectors represent Fourier modes and are used for defining the GFT. The GFT of brain signal 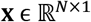 is obtained as

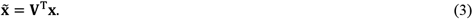

The inverse GFT of 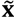 is attained by

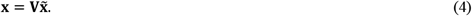

It is noteworthy that the eigenvector corresponding to the larger eigenvalue shows more variance and can be regarded as the graph higher frequency (Huang et al., 2018). The graph signal can be transformed to the graph frequency domain, and then filtered at that domain, and finally got inversed process using “(4)” in order to have graph filtered signal. This filtering process is expressed as

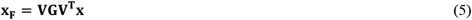

where **G** is a diagonal filtering matrix. In this study, we consider 1 for the diagonal elements corresponding to the desired frequency modes and 0 for the rest modes.

### 2.4. Scenarios

In this study, the graph frequency modes are divide into 8 separate bands and applied to the rfMRI data using graph filtering. The number of frequency modes contributing in first to eighth bands are 1-12, 13-24, 25-36, 37-48, 49-60, 61-72, 73-84, and 85-97, respectively. As with (Huang et al., 2018), we consider a few number (close to 10) of frequency modes for each band. Hereinafter, our means of frequency is graph frequency unless otherwise stated.

In this study, the first and eighth frequency bands are respectively considered as low and high frequency bands. In low frequencies, the physically connected ROIs activate similar to each other, whereas in the high frequencies, the variance of ROIs activities with respect to structural graph is high.

In the following of this subsection, different measures and their computation methods are explained. These measures relation to age and also their diagnosing powers are studied in this paper.

#### 2.4.1. MCC analysis

In this scenario, a CC analysis similar to (Huang et al., 2018) is carried out. The CC is computed as the root mean square of rfMRI filtered data at each time point, and then is averaged across total time points of a given subject in order to obtain mean CC (MCC). The MCC is computed for each of 7 understudy networks and also for total brain.

#### 2.4.2. QCC analysis

In quartile CC (QCC) scenario, the CC and also the first, second, seventh, and eighth frequency bands are used for total brain analysis. At each band, each CC value is replaced by quartile this CC falls in based on total CC values of understudy band for a given subject. Thus, for each subject and each understudy band, there exists a time series of signal with values 1, 2, 3, and 4. At each time point, the quartile values of four understudy bands form a vector called meta concentration coefficient (*meta-CC*) (Mennigen et al., 2018).

For the first analysis, similar to (Mennigen et al., 2018), four global metrics of CC level changes are examined:

1. The number of *meta_CC* changes for each subject (represented by *s*).
2. The number of unique *meta_CC* for each subject (represented by *n*).
3. The largest Euclidean distance between two *meta_CCs* for each subject (represented by *r*).
4. The sum of distance between consecutive *meta_CCs* for each subject (represented by *d*).

For the second analysis, in the first and eighth bands, the CC level transition probability is computed by a first order hidden Markov model for each subject. This analysis expresses the probability of entering into a new level or staying at the current level.

Some other analyses were performed and significant results were not obtained. These analyses and their corresponding results have been reported in section S.2 of supplementary file.

#### 2.4.3. FC analysis

In this scenario, the FC between ROIs and also within and between networks are investigated in low and high frequency bands. To this end, the FC is computed between ROIs using Pearson correlation, and then is z-scored. For considering the transient nature of brain FC, the SW technique is employed for FC calculation. In SW, a sequence of widows with one TR shift are applied to each ROI time series, and then one FC matrix is computed for each window. The final correlation value of a ROI-ROI would be the mean of that ROI-ROI FC values. The window is created by convolving a rectangle (width = 22 TRs) with a Gaussian (σ = 3 TRs) (Allen et al., 2014).

For obtaining each within network (e.g., DMN-DMN) and each between networks (e.g., DMN-SCN) FC values, their corresponding ROI-ROI z-scores are averaged. Thus, it is totally attained 7 within network and 21 between networks FC values for a given subject.

### 2.4.4. Cross frequency FC analysis

In this scenario, the FC is computed between ROIs low frequency signals and high frequency signals in order to compute cross frequency FC matrix. This matrix is not symmetric and each element of which is the correlation value existing between one ROI low frequency signal and that or another ROI high frequency signal. The implementation of this scenario is similar to FC analysis one and the final ROI-ROI and within and between networks FC values are calculated the same as explained for FC analysis scenario. In this study, each row/column of cross frequency FC matrix represents the FC values existing between low/high frequency signal of a given ROI and high/low frequency signals of 97 understudy ROIs. Hereinafter, the ROIs with low/high frequency signals are stated as low/high frequency ROIs and the networks with low/high frequency ROIs are stated as low/high frequency networks.

### 2.4.5. LVFS analysis

The short time frequency transform (STFT) has been done by discrete Fourier transform of time series signal and a spectral domain window. Similar approach can be performed for graph signal in graph spectral domain in order to localize a vertex in graph spectral domain (Stankovic et al., 2019). The LVFS for a vertex *m* and spectral index *k* is computed by

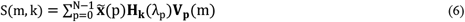

where **V**_p_ is p^th^ graph frequency mode and **H**_*k*_ is a band-pass transfer function. In this study, 8 **H**_*k*_ are used which are corresponding to 8 understudy frequency bands. These transfer functions are illustrated in Fig 1. The LVFS (S) is computed for each vertex and transfer function so that one matrix with dimension 8 × 97 (the number of frequency bands × the number of vertices) is derived at each time point for a given subject. In order to reduce the LVFS susceptibility to random variations originating from noise artifacts and also because of computing LVFS for different subjects, the LVFSs are z-scored. To do this, the graph signal at each time point is shuffled 200 times and consequently 200 different LVFS matrices are computed. Then, each S(m, k) of original LVFS matrix is z-scored. For each time point, this z-scored process is carried out. The z-scored LVFSs of ASD and TC subjects are separately clustered using k-means with 400 times repetition and a maximum iteration of 200. The Elbow metric which is a ratio of within cluster sum of squared distances (WCSSD) to between clusters SSD (BCSSD) is employed for finding the optimum number of clusters. For WCSSD, for a given cluster, the SSD of all members of that cluster from the center of that cluster is computed and this process is repeated for all clusters. Finally, the sum of all of these SSDs is considered as WCSSD. For BCSSD, for a given cluster, the SSD of all LVFSs which are not a member of that cluster from the center of that cluster is computed and this process is repeated for all clusters. Finally, the sum of all of these SSDs is considered as BCSSD. After clustering, the clusters’ centers and members respectively represent the LVFS’ states and states’ members. This analysis is similar to DFC one with one difference which is using LVFS matrices instead of FC ones.

**Fig 1.**
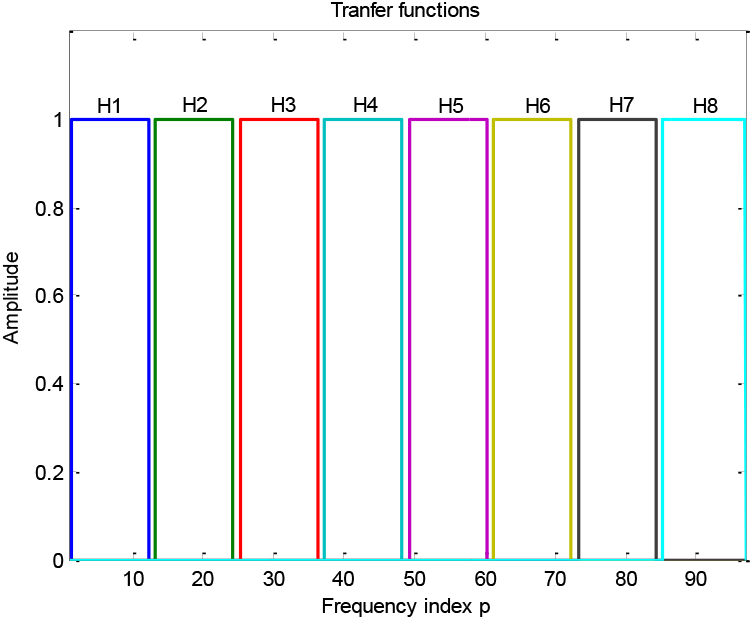
Eight transfer functions in the spectral domain.

For each ASD state, the closest TC state to that is specified using Euclidean distance so that the final matched states are the best matching pairs. In each state, all LVFS matrices belonging to a subject are averaged to have one LVFS matrix for that subject. Then, in each state, the LVFS value differences between ASD and TC is analyzed at vertex and network level for a given spectral band. For network level, the LVFS of each network is computed by averaging the LVFSs of its ROIs.

## 2.5. Statistical analysis

### 2.5.1. Analysis the age and diagnosis variables effects on measures

As it was explained in scenarios subsection, many measures are investigated in this paper. For both ASD and TC, the SDSU data mostly includes males and so the reliable statistical analysis of sex effects is not possible. Therefore, only the effects of age and diagnosis variables on measures are examined. The diagnosis variable **x**_diagnosis_ is binary, with ASD coded as ‘1’ and TC as ‘0’. This variable determines which measure can be a significant discriminative feature between ASD and TC.

The first linear model is for analyzing the age variable effect and is defined as

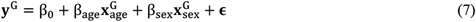

where G represents ASD or TC group and Y is measure vector. The second linear model is for analyzing the diagnosis variable effect and is defined as

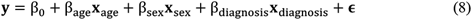

where the **y** is a vector containing both ASD and TC measures, so β_diagnosis_ > 0 indicates a positive correlation with ASD and β_diagnosis_ < 0 shows a negative correlation with ASD. The β_diagnosis_ with a *p* < 0.05 indicates that the understudy measure is significantly different between ASD and TC groups.

These linear models are performed using the command “stepwiseglm” of MATLAB software with the constant model as the starting model, linear model as the upper model, and deviance as the criterion for adding or removing terms. If understudy variable does not exist in the general linear model created by stepwiseglm, the command “addterms” is used in order to add the understudy variable to the created model and consequently the *β* and *p* of that variable are obtained. This adding process is implemented because the *β* and *p* of understudy variable are needed to multiple comparison correction.

### 2.5.2. Multiple comparison correction

For each ASD or TC group, many *β* and corresponding *p* values are computed for each age and diagnosis effects. For example, at the first scenario, 7 networks and total brain in low and high graph frequency bands form 16 situations and analysis of these situations results in 16 *β* and 16 *p* values. For other scenarios, a lot more situations exist. Consequently, multiple comparison problem is occurred. In this study, the pixel-based multiple comparison correction (PBMCC) is exploited in which each situation presents one pixel (Cohen, 2014). In PBMCC technique, the permutation testing is iterated *Q* times and in each time the largest positive and the largest negative pixel values are saved. The pixel value in this study is –*sign*(*β*)*log*_10_(*p*). After all iterations have completed, two distributions of the largest positive and the largest negative pixel values are attained. For situations with original –*sign*(*β*)*log*_10_(*p*) > 0 and –*sign*(*β*)*log*_10_(*p*) < 0, respectively, the distribution of the largest positive and the distribution of the largest negative values are employed in order to verify or reject the significance of their *p-values*.

Two different permutation testing are used for 5 scenarios measures. For the first two scenarios, the Fourier phase randomization procedure (Huang et al., 2018; Theiler et al., 1992) and then graph phase randomization (Huang et al., 2018) are applied to the graph signals. The first randomization which is in temporal dimension can be expressed as

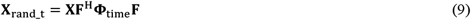

where.^H^ indicates the Hermitian transpose, **F** is the Fourier matrix, and the diagonal of **Φ**_*time*_ contains random phase factors. The second randomization which is in spatial dimension is defined as

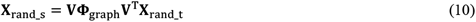

where the random sign flips are stored on the diagonal of **Φ**_*graph*_. In this randomization, the graph spectral coefficients are randomized by flipping their signs. More information about these randomizations can be found in (Huang et al., 2018). By these randomizations, different signals are generated for ASD and TC and then two positive and negative distributions are formed for each measure. Each of these randomizations may be enough but we are interested in employing both of them.

The fifth scenario deals with clustering process and above mentioned randomizations are possible to change the optimized cluster numbers. When the permuted cluster numbers become different from the original ones, the multiple comparison correction would not be possible. Therefore, for measures of this scenario in each iteration of permutation testing, we shuffle the measure vectors ***y*** and **y**^G^ while keeping untouched the predictor variables 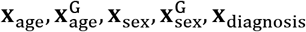. By shuffling *𝒢* times, the two positive and negative distributions are formed.

It is known that phase randomization preserves the correlation among signals. Therefore, for the third and fourth scenarios in which the FC is computed using Pearson correlation, the phase randomization cannot be exploited for permutation testing. Hence, as with the fifth scenario, the shuffling techniques is used.

In this study, the *Q = 300* and the *p-value* correction of age variable is carried out for ASD and TC groups separately. In all obtained distributions, the null hypothesis is that there is no significant difference between ASD and TC (for the second linear model) or is that there is no significant relation between understudy measure and age variable (for the first linear model).

## 3. Results

### 3.1. MCC results

Two groups didn’t show significant difference in term of MCC in both low and high frequencies (p > 0.3 for all 7 networks and total brain). For TC group, the MCC was independent from age (p > 0.3 for all 7 networks and total brain). The MCC for ASD group showed significant relation to age for networks including DMN, CCN, SMN, VN, CN, SCN and also the total brain (all *p* < 0.01; all *β* < −1) in low frequency band. In high frequency band, the age significantly impacted on networks DMN, CCN, VN, CN and SCN in term of MCC (all p < 0.04; all *β* < −1).

### 3.2. QCC results

For the first analysis, only the fourth metric *d* was significantly different between ASD and TC groups (p < 0.05; *β* = −20.17). For age variable, the ASD group showed significant positive relation only to fourth metric (p < 0.05; *β* = 4.37).

For the second analysis, the transition probabilities of ASD and TC were separately averaged over subjects in order to obtain the mean transition probability. The differences between means of ASD and TC in low and high frequencies are shown in Fig 2. The probabilities of transition from CC levels 1,1,4,4, respectively, to 1,3,1,3 were significant for diagnosis variable in low frequency band. In high frequency band, the probability of transition from CC level 4 to 1 was significant. It wasn’t seen any significant relation to age.

**Fig 2.**
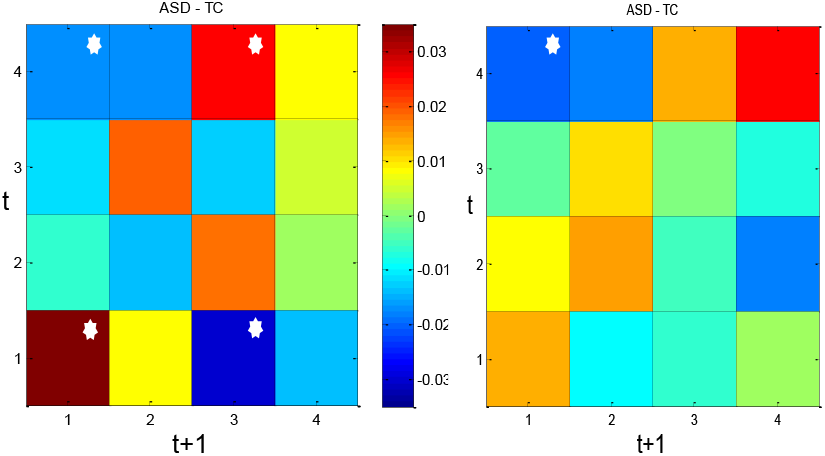
Mean of CC level transition probability of ASD subtracted from that of TC for low frequency (left) and high frequency (right) bands. The *t* denotes time and transitions showing significant difference between ASD and TC are indicated by a white star (*p* < 0.05).

### 3.3. FC results

The results of FC analyses at ROI and network levels are shown in Fig 3. In low frequency band, for ASD, the significant negative effect of age variable on most of the ROI-ROI FCs, except those of contributing in DMN-DMN, CN-SCN, and AN-AN, can be seen. Some ROI-ROI FCs of CCN-VN show significant positive relations to age. In contrast, the significant positive relation of ROI-ROI FCs to age is dominant for TC group. These relations mostly exist in DMN-DMN, CN-SCN, SMN-CN, CCN-CN, and AN-CN. For diagnosis variable, the most ROI-ROI FCs particularly those contributing in CN-SCN, VN-CN, CCN-CN, DMN-CN, and DMN-CCN are significantly more positive or less negative for ASD in comparison to TC. In high frequency band, no ROI-ROI FC can survive from multiple comparison correction.

**Fig 3.**
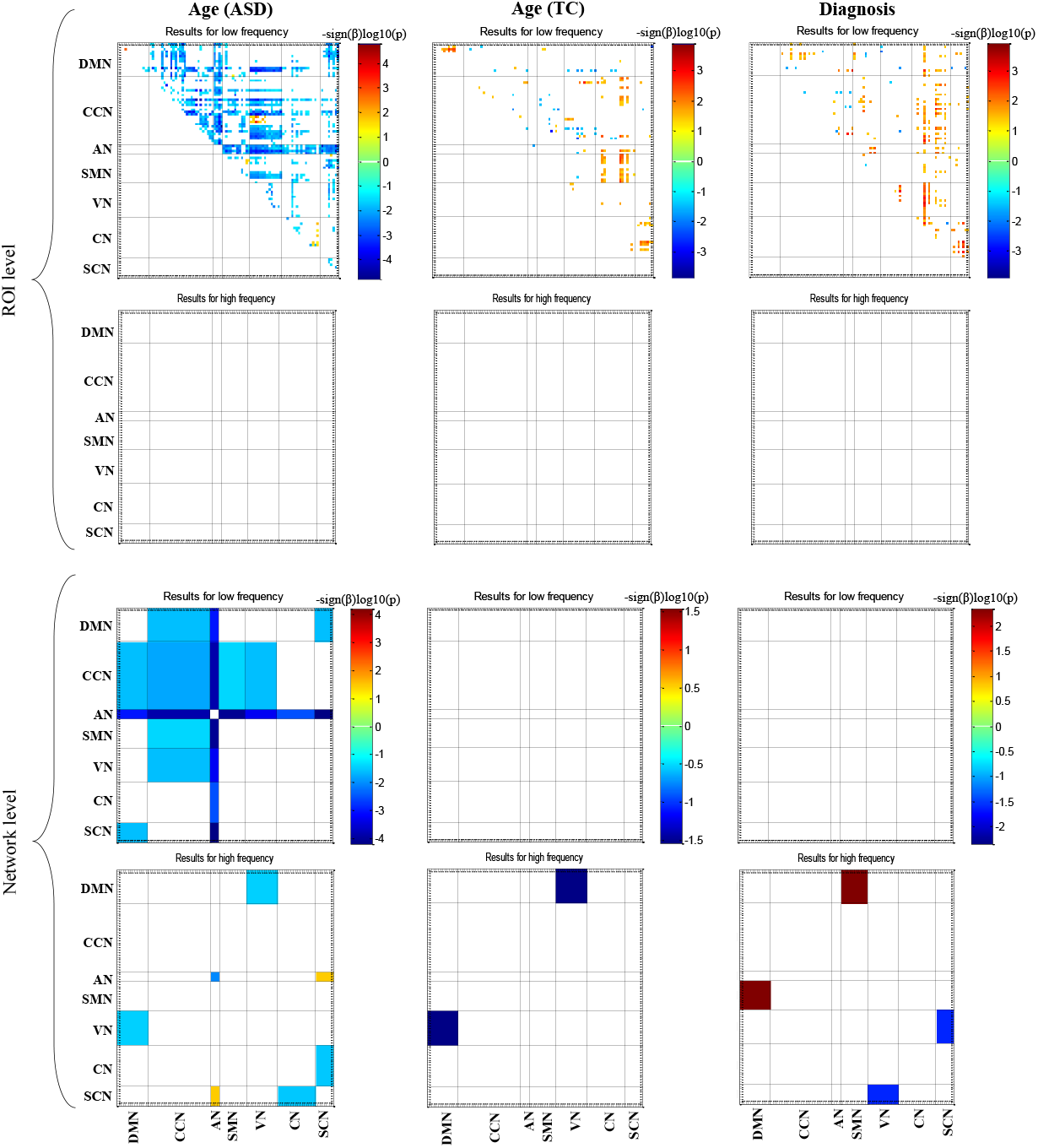
Functional connectivity analyses with respect to age and diagnosis variables in low and high frequency bands. The results are reported at ROI and network levels. Significant results (*p* < 0.05) are shown by color related to –*sign*(*β*)*log*10(*p*).

For network level analysis, in low frequency band, the age effect on between and within networks FCs of ASD is only negative. Particularly, the AN connections with all networks except itself is obtaining less positive or more negative by increasing the age. The CCN shows similar behavior to AN but with less strength and only with DMN, CCN, AN, SMN, and VN. The connections of DMN with CCN, AN, and SCN also represent anti-correlation behavior with age. The age variable for TC group and the diagnosis variable do not show any significant relations to between and within networks FCs.

In high frequency band, the AN-AN and AN-SCN relations to age are negative and positive, respectively. Both ASD and TC groups represent significantly negative relations to age for DMN-VN. For diagnosis variable, the ASD/TC has more correlation or less anti-correlation than TC/ASD for DMN-VN/VN-SCN. In contrast to ROI level, the useful results of network level analysis are in high frequency band.

The network level analyses of diagnosis variable are also implemented in six remained frequency bands. The results of this analyses are shown in section S.3 of supplementary file. Both positive and negative relations to diagnosis variable exist in these bands, particularly in the middle and higher frequency bands. Some within and between networks FCs keep their relation signs to diagnosis variable in different bands, ex., the DMN-VN while some others change their relation signs ex., CCN-VN.

### 3.4. Cross frequency FC results

The results of this scenario for age and diagnosis variables are shown in Fig 4. The cross frequency FC matrices of ASD and TC are shown in Fig. S5 of supplementary file. In this scenario, in addition to ROI-ROI and network-network analyses, a network-ROI analysis is also performed. In network-ROI, for a given low/high frequency ROI, its FC values with ROIs of a given high/low frequency network are computed and then averaged. By doing so, it is determined that how much a given low/high frequency ROI is connected to the given high/low frequency network. Also, this analysis can show the different behaviors of a ROI in low and high frequency bands.

**Fig 4.**
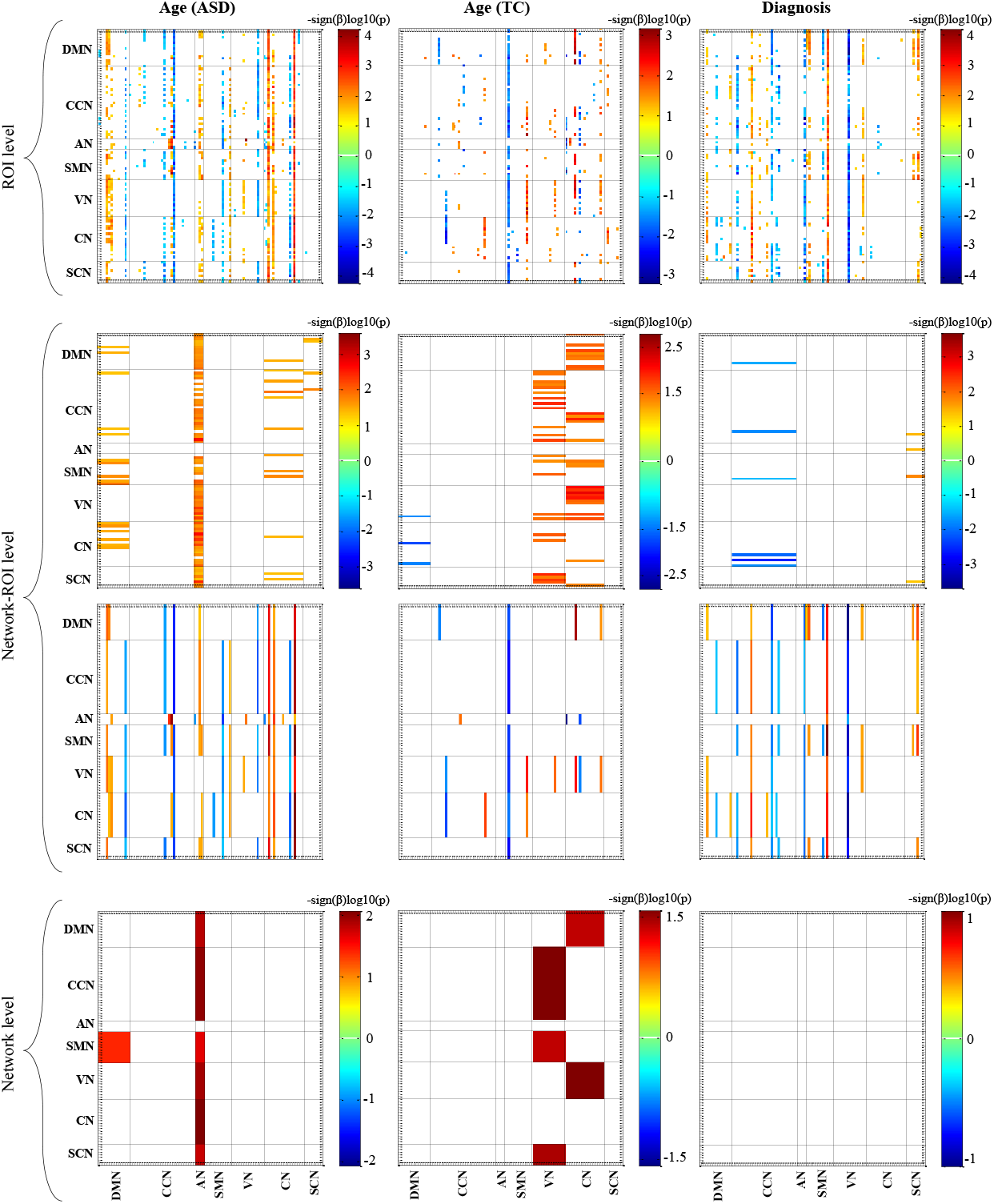
Cross frequency FC results (–*sign*(*β*)*log*10(*p*)) for age and diagnosis variables at ROI, network-ROI, and network levels. Only the significant relations to age and diagnosis variables (*p* < 0.05) are shown.

The column/row view is related to FC values existing between a given high/low frequency ROI and all low/high frequency ROIs. At ROI level, the column view offers regular patterns across low frequency ROIs. The correlations of some high frequency ROIs with most of the low frequency ROIs are significantly related to age variable. For ASD, these some ROIs which represent the positive relations to age are the left and right anterior Cingulum and left middle Cingulum (ROIs of DMN), left and right superior temporal (ROIs of AN), right paracentral lobule (ROI of SMN), cerebellum 3 and left cerebellum Crus 2 and particularly left cerebellum 9 (ROIs of CN). The negative relations are dominant for right medial superior frontal (ROI of DMN), right parahippocampal and insula (ROIs of CCN), left supramarginal (ROI of SMN), righ inferior occipital (ROI of VN) and left cerebelum 8. For TC, less column wise regular patterns exist. The most negative relations to age are for right precentral (ROI of SMN) and the most positive relations are for right supramarginal (ROI of SMN), left cerebelum 3, and left cerebelum 10 (ROIs of CN).

For diagnosis variable, some high frequency ROIs show regular patterns of positive or negative diagnosis effects approximately across all low frequency ROIs. These high frequency ROIs are right pallidum (SCN, “+”), left superior occipital (VN, “-”), right supramarginal (SMN, “+”), right superior temporal (AN, “-”), right hippocampus (CCN, “-”), opercular part of left inferior frontal (CCN, “+”), orbital part of left superior frontal (CCN, “-”), orbital part of right medial frontal (DMN, “+”). The information inside the parentheses respectively are the network and dominant diagnosis effect sign of given ROI. The positive/negative effect for a high frequency ROI means that that ROI correlations with low frequency ROIs for ASD/TC is more positive or less negative than TC/ASD. The FC values related to these high frequency ROIs for ASD and TC are shown in Fig. S6. The left superior occipital and right hippocampus show dramatic difference between ASD and TC so that the TC FC values are positive and those of ASD are negative. For right superior temporal, the ASD FC values are negative but at least half of the TC FC values are close to zero and the rest are positive. The opposite dramatic difference between ASD and TC can be seen for right supramarginal and opercular part of left inferior frontal where the FC values of ASD are positive and those of TC are negative. Such opposite behavior can also be seen for right pallidum where TC FC values are negative and those of ASD are positive. However, it should be mentioned that most of the ASD FC values of right pallidum are close to zero. For orbital part of left superior frontal/orbital part of right medial frontal, the FC values of TC/ASD are much less/less negative than those of ASD/TC. Also, two other high frequency ROIs which are left fusiform (VN, “+”) and right precentral (SMN, “+”) attract the attention. For these ROIs, the ASD FC values are positive (close to zeros for a few ROIs) and TC FC values are negative and somewhat close to zero. The most interesting result is that the differences between ASD and TC are observed across approximately all of the low frequency ROIs. However, these difference across some low frequency ROIs are much more.

The network-ROI level analysis can show these differences in a much more localized view. Also, the network-ROI level analysis enables finding the ROIs which can be highly correlated with one network when they are only in high or only in low frequency band. The second and third rows of Fig 4 show the FCs having the significant relations to age and diagnosis variables. The FCs in second/third rows of this figure are the connections between one given low/high frequency ROI and high/low frequency networks. Interestingly, in both rows of this figure, regular patterns are seen from column view. For second row of Fig 4, this means that the connections of some high frequency networks with the most low frequency ROIs have significant relations to age variable and also are significantly different between ASD and TC. For third row of Fig 4, this means that the connections of some high frequency ROIs with most of the low frequency networks have significant relations to age and diagnosis variables. Overall, based on this scenario results, it can be stated that it is only probable for high frequency band that some of its ROIs or networks connect to most of the low frequency ROIs so that these connections offer significant difference between ASD and TC and significant relations to age.

For age variable results of ASD, the connections of high frequency DMN, CN, and particularly AN with most of the low frequency ROIs offer significant positive relation. Also, the connections of high frequency ROIs including the left cerebellum 9, left cerebellum 3, and left cerebellum crus 2 (CN) and right parahippocampal (CCN) with most of the low frequency networks are significant in relation to age. For TC, high frequency VN and CN and high frequency right precentral (SMN) offer significant relations to age variable. For diagnosis variable, the high frequency CCN connections with some low frequency ROIs are significantly less for ASD in comparison to TC. The high frequency ROIs reported for ROI level results again attract the attention at network-ROI level. The connections of these high frequency ROIs, particularly left superior occipital (VN, “-”) and right supramarginal (SMN, “+”), with all or most of low frequency networks are significantly different between ASD and TC. The connections related to left superior occipital/right supramarginal are negative/positive for ASD and positive/negative for TC.

At network level, only significant positive effects are found for age variable. For ASD, the positive effects of age can be seen for connections existing between high frequency AN and the rest networks when they are in low frequency band. Also, the positive effect on connection between low frequency SMN and high frequency DMN exists. For TC, by increasing/decreasing the age, the connections between low frequency VN and high frequency CCN, SMN, SCN are increased/decreased. The similar effect of age is seen for connections between high frequency CN and low frequency DMN, VN. The diagnosis variable doesn’t show any significant difference between ASD and TC at network level.

### 3.5. LVFS results

The LVFS states of ASD and TC groups are shown in Fig 5. The optimum cluster number for both groups is 5 (see section 4 of supplementary file). Both groups show similar local vertex spectrum behavior in all 5 states and the large positive and negative LVFS values exist in the second and particularly in the first graph frequency bands (GFB1 and GFB2). In states 1 to 4, dominantly the LVFS behaviors of VN and DMN and somewhat of SMN attract the attention. In states 1 and 4, the vertices of VN respectively have the largest negative and positive S whereas in states 2 and 3, the vertices of DMN have the largest negative and positive S, respectively.

**Fig 5.**
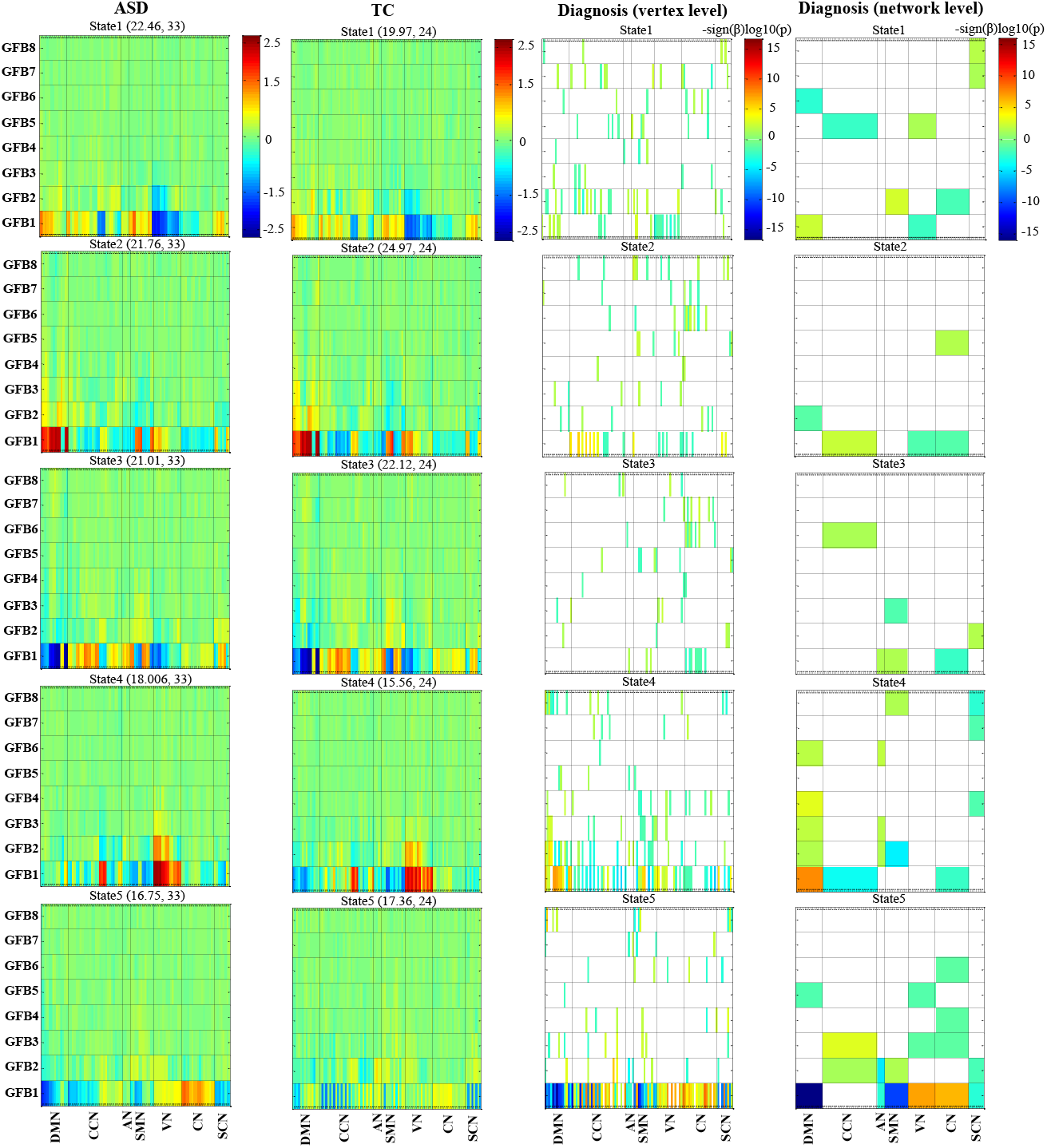
The first and second columns (from left to right) represent the ASD and TC LVFS states. Each state contains frequency spectrum information of 97 vertices organized in 7 brain networks at 8 different and completely separate graph frequency bands (GFBs). The numbers above each state from left to right respectively express the percentage of time points contributing in that state and the number of subjects who have at least one time point at that state. The third and fourth columns represent the diagnosis variable analysis results (–*sign*(*β*)*log*10(*p*)) at vertex and network levels. Only the significant relations to diagnosis variable with *p* < 0.05 are shown.

All subjects usually do not experience each DFC state (Mash et al., 2019; Rashid et al., 2018) whereas all subjects have contributions in each LVFS state. In states 1 and 4, a significantly larger proportion of ASD than TC time points contribute (**state1**: ASD = 22.46%, TC = 19.97%; χ^2^ (1) = 7.94, < 0.005 ::: **state4**: ASD = 18.006%, TC = 15.56%; χ^2^ (1) = 9.11, *p* < 0.005). In state2, the proportion of TC time points is significantly larger than the TC ones (**state2**: ASD = 21.76%, TC = 24.97%; χ^2^ (1) = 12.52, *p* < 0.0005).

The significant results of diagnosis variable (p < 0.05) at vertex and network levels are shown in Fig 5. The most significant differences between ASD and TC at both vertex and network levels are found in state4 and particularly in state5 and at the two lowest frequency bands including GFB1 and GFB2. In state4 and GFB1, the TC group has negative LVFS values for roughly half of the DMN vertices whereas the ASD group has zeros or positive LVFS values for DMN vertices. Thus, significant positive effect of diagnosis variable is reported for DMN and more of its vertices. The similar behavior but this time for TC with respect to ASD can be seen for DMN and its vertices in state5. As a result, significant negative effect of diagnosis variable is seen in this state for DMN and its vertices. The SMN vertices of TC have less negative or more positive frequency spectrum values in comparison to those of ASD in GFB1. Consequently, negative diagnosis variable effects are seen for SMN and its vertices. The VN and CN vertices of ASD respectively have positive and highly positive frequency spectrum values localized in GFB1 whereas those values of TC are zero or low positive. Therefore, the VN and CN and their vertices show positive effects for diagnosis variable. For age variable, no significant effect is observed for both ASD and TC at vertex level. At network level, negative effects for ASD are observed in state1 (DMN at GFB1) and state3 (SCN at GFB8). For TC, only frequency spectrum values of SCN (at GFB8) in state3 has negative relation to age.

## 4. Discussion

Herein, the SDSU dataset of ABIDE II database was used in order to analyze the rfMRI data behavior in topological domain through GFT. For topological domain, the underlying structure of rfMRI data was considered the structural connectivity attained by DTI data. Various scenarios in relation to signal concentration, FC, and LVFS were performed in different graph frequency bands.

Based on the MCC scenario results, it can be said that the signal concentration of most of the brain networks are decreased by increasing the age for ASD group.

For the QCC scenario, the results of second analysis show that if ASD subjects are at CC level one/four in low frequency band, their intension for staying in that level or going only one level higher/lower is more than TC subjects. In high frequency band, such an intention can be seen for level four. In contrast, the TC subjects in comparison to ASD ones have more intentions for transiting from level four to much lower level in both low and high frequency bands and also from level one to much higher level in low frequency band. These findings are consistent with the result of first analysis where the sum of distance between consecutive *meta_CCs* (denoted by *d*) was significantly less for ASD in comparison to TC.

Based on the FC analyses at ROI level and in low frequency band, it can be said that increasing the age results in increasing the segregation/integration of ASD/TC subjects brain ROIs. By removing the age variable effect for diagnosis variable, it was found overall more integration of ASD than TC in low frequency band. For both age and diagnosis variables, no ROI level FCs survived from multiple comparison correction in high frequency band. Increasing the segregation of brain networks particularly AN, SCN, and DMN by increasing the ASD subjects’ age was another founding of this research in low frequency band. In contrast to ROI level, the most significant information of network level analyses was in high frequency band. At this band, increasing the integration of ASD subjects’ AN and increasing the segregation of DMN for both ASD and TC were results of increasing the age. The findings related to AN expressed the fact that one brain network functional behavior can be significantly different in different graph frequency bands. Increasing the segregation of ASD subjects’ DMN by increasing the age was another typical findings. By removing the age variable effect, the diagnosis variable revealed the reduced segregation of ASD subjects’ DMN (reduced anti-correlations or increased positive connectivity with SMN) (Mash et al., 2019).

The network level analysis of diagnosis variable in frequency bands second, third, fourth, fifth, sixth, seventh and eights demonstrated the fact that many distinctions between ASD and TC in term of FC can be obtained from middle and higher frequency bands. Therefore, the belief that only the first several graph frequency modes contain the most significant information just as the classical Fourier frequency (Wang et al., 2018) cannot be valid at least for comparison analysis of rfMRI data of ASD with respect to TC. Some between networks FCs including DMN-VN, SMN-VN and CN-CN keep their sign of relation to diagnosis variable at different frequency bands whereas some others including SMN-SCN and CCN-VN change their signs. From these between networks behaviors it can be stated that each frequency band may reveal one aspect of between network information which can be band specific information. Also, these behaviors may be a way for justifying some of the inconsistent findings of SFC in which it is possible some researches find positive signs and some others report the negative signs. The justification may be that each finding is represented in a specific band. This manner of justification means that some inconsistent findings may be justified by applying more detailed analyses such as investigating the SFC in different graph frequency bands. Taking such detailed investigation into account may provide results which are consistent across different studies. Recently, it has been stated that the DFC technique can be a method for answering to the inconsistent findings of SFC by attaining those findings in different states (Mash et al., 2019). The GFT may be another approach for answering to these findings.

For the cross frequency FC scenario, the low and high frequencies were defined in topological domain and the FCs were computed in temporal domain. At ROI and network-ROI and network level analyses, only the results related to high frequency band offered regular patterns. This means that only some high frequency ROIs and networks were found that their connections with the most low frequency ROIs or networks had significant relations to age and diagnosis variables. In contrast, the low frequency ROIs and networks connections with a few high frequency ROIs or networks had significant relations to age or diagnosis variables. These few numbers were not enough for forming a regular pattern. In previous scenarios, the ASD and TC usually offered close values for the calculated statistically significant measures, meaning that both groups had positive or negative values for the given measure or one had negative/positive and the other had less negative/positive values. In this scenario, the connections of high frequency ROIs including left superioroccipital, right hippocampus, right supramarginal, and opercular part of left inferior frontal with the most low frequency ROIs were dramatically and significantly different between ASD and TC. One group had the positive FC values and the other one attained negative FC values. Hence, the FC analysis of rfMRI data between graph low and high frequency bands may can provide powerful tools for introducing candidate biomarkers. Overall, cross frequency FC analysis can be a new avenue for comparison study between at least ASD and TC groups.

In LVFS scenario, the vertices spectrum were localized in 8 different frequency bands. The most important information of LVFS matrices of a lot of time points and subjects were provided by clustering and consequently the state analysis tools. The percentage of time ASD subjects spent in states 1 and 4 were significantly more than TC subjects and in these states the LVFS values of VN vertices were dominant in GFB1. In contrast, the TC subjects devoted more percentage of their times than ASD subjects in state2 where the DMN vertices had the largest positive spectrum values in GFB1. In GFB1 of state5, the ASD subjects had much stronger LVFS values than TC subjects in VN and CN. The DMN vertices of ASD and TC subjects in GFB 1 showed contradictory behavior in states 4 and 5. The values of many of these vertices in state4/state5 were positive/negative for ASD and negative/positive for TC. By considering the age variable results reported in section 3.5, it may can be said that the most significant results reported for diagnosis variable are not sensitive to age. Overall, it can be said the LVFS analysis at different states can offer valuable information about differences existing between ASD and TC. For this scenario, it can be stated that the most significant information of LVFS is found at graph low frequency bands (GFB2 and particularly GFB1).

Another way of analysis is employing tensor factorization technique for ASD and TC data separately (Kolda and Bader, 2009). A 3D tensor can be employed where the first, second and third dimensions stand for FC vector (the upper triangular of FC matrix is represented as vector), subjects, and graph frequencies, respectively. This scenario has been carried out and for interested readers the results have been reported in section 7 of supplementary file.

### 4.1. Limitations and future directions

The limitations of this study are related to the used data. One of limitations of this work was the number of subjects. Using much more subjects can more reliably extend the results of this study to the ASD populations. Another source of limitations was the broad range of understudy subjects’ age which was from 7.4 to 18 years. The number of females was very limited and so the sex effect on understudy measures was not investigated.

For future works, the measures of this paper can be applied to the rfMRI and DTI data with a lot more subjects and females and for a narrow range of year. In this study, the different graph frequency bands were investigated when the temporal dimension Fourier frequency was 0.01-0.1 Hz. However, it is possible to investigate many measures of this paper in sub-bands of 0.01-0.1 Hz. By doing so, both graph and temporal frequencies are analyzed simultaneously. Another work for future can be applying the DFC in graph low and high frequency bands and consequently analyzing its differences and commonalities in these bands. The QCC analyses can be employed to FC (QFC). For QFC, many FC values of desired ROI-ROIs can be computed by employing the sliding window technique for a given subject. Final suggestion is using GSP for task fMRI data of ASD and TC.

## 5. Conclusion

In this paper, the rfMRI data of ASD and TC subjects were analyzed using GSP. The analyses were performed in different graph frequency bands and different scenarios. The MCC and QCC analysis in low and high frequency bands could offer new knowledge about signal concentration of ASD and TC and also differences existing between them. Increasing the age leaded to increasing segregation of ASD subjects’ brain at both ROI and network levels in low frequency band. Reduced segregation of DMN in ASD occurred in middle and higher frequency bands. The cross frequency FC analysis, performed between low and high frequency signals, showed that some high frequency ROIs can provide significant diagnostic features. The left superior occipital, right hippocampus, right supramarginal and opercular part of left inferior frontal regions in high frequency provided dramatic significant different FC values for ASD and TC. The results of LVFS analysis demonstrated that investigating the LVFS of brain networks in different states can be a good idea for comparing the ASD and TC in aspect of their localized frequency spectrum values. The results of all scenarios confirm that using both structural and functional data through GSP tools can open a new reliable avenue for recognizing the ASD.

## Supporting information

Supplementary file of paper

---

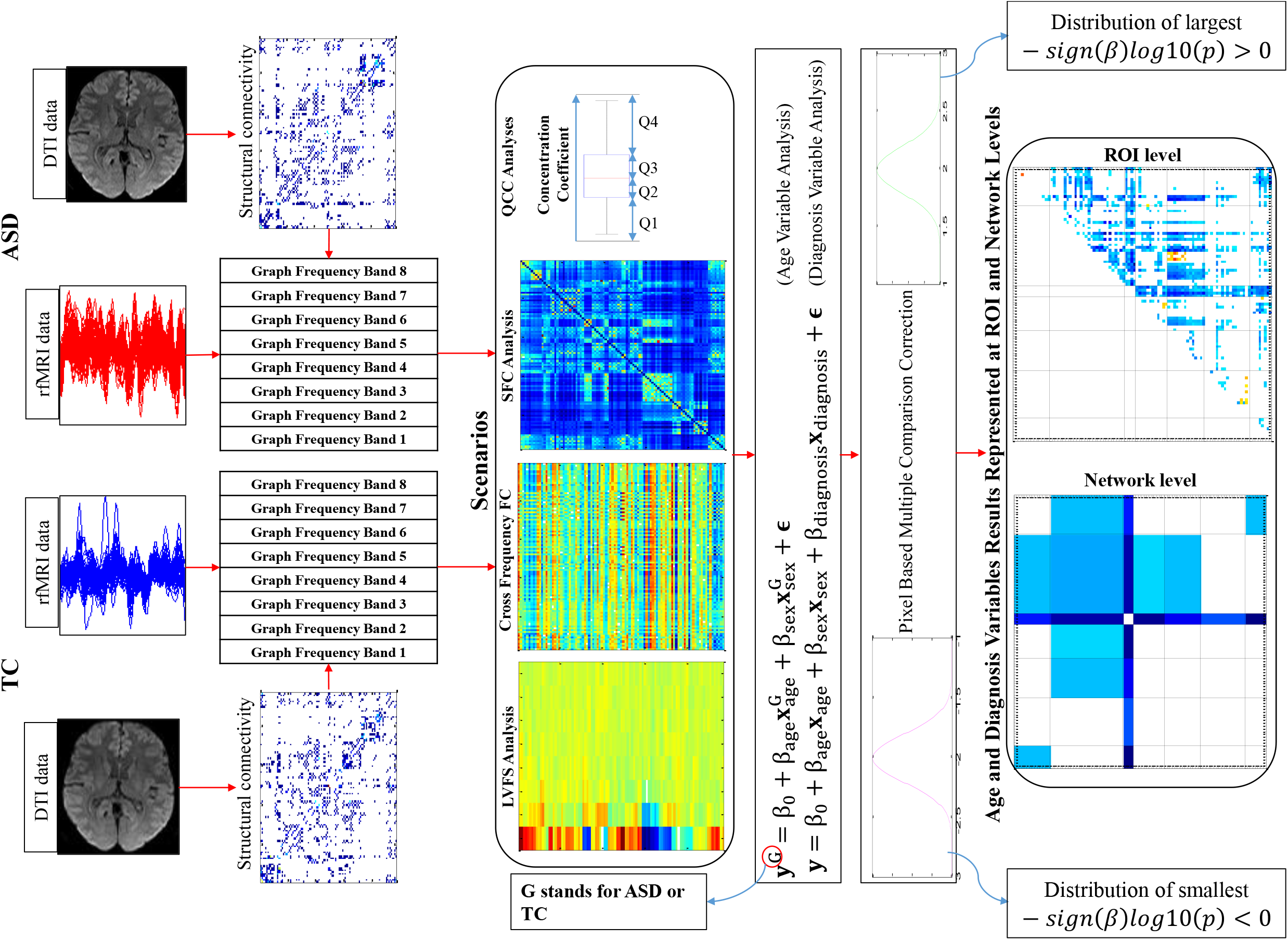

1 http://www.fil.ion.ucl.ac.uk/spm/.

2 https://afni.nimh.nih.gov.

3 http://fmri.wfubmc.edu/software/pickatlas.

## References

Aggarwal, P., Gupta, A., 2019. Multivariate graph learning for detecting aberrant connectivity of dynamic brain networks in autism. Medical image analysis, 56, 11–25.

Allen, E. A., Damaraju, E., Plis, S. M., Erhardt, E. B., Eichele, T., Calhoun, V. D., 2014. Tracking whole-brain connectivity dynamics in the resting state. Cerebral Cortex, 24(3), 663–676.

American Psychiatric Association, 2013, Diagnostic and statistical manual of mental disorders. American Psychiatric Association, Washington DC.

Biswal, B.B., 2012. Resting state fMRI: a personal history. Neuroimage 62, 938–944.

Brahim, A., Farrugia, N., 2020. Graph Fourier Transform of fMRI temporal signals based on an averaged structural connectome for the classification of neuroimaging. Artificial Intelligence in Medicine, 101870.

Buescher, A. V., Cidav, Z., Knapp, M., Mandell, D. S., 2014. Costs of autism spectrum disorders in the United Kingdom and the United States. JAMA pediatrics, 168(8), 721–728.

Caballero-Gaudes, C., Reynolds, R. C., 2017. Methods for cleaning the BOLD fMRI signal. Neuroimage, 154, 128–149.

Cheng, W., Rolls, E. T., Gu, H., Zhang, J., Feng, J., 2015. Autism: reduced connectivity between cortical areas involved in face expression, theory of mind, and the sense of self. Brain, 138(5), 1382–1393.

Cohen, M. X., 2014. Analyzing neural time series data: theory and practice. MIT press.

Di Martino, A., Yan, C.-G., Li, Q., Denio, E., Castellanos, F. X., Alaerts, K., Anderson, J. S., Assaf, M., Bookheimer, S. Y., Dapretto, M. et al., 2014. The autism brain imaging data exchange: towards large-scale evaluation of the intrinsic brain architecture in autism. Molecular psychiatry, 19, 659.

Di Martino, A., O’connor, D., Chen, B., Alaerts, K., Anderson, J. S., Assaf, M. Blanken, L. M., 2017. Enhancing studies of the connectome in autism using the autism brain imaging data exchange II. Scientific data, 4(1), 1–15.

Drysdale, A. T., Grosenick, L., Downar, J., Dunlop, K., Mansouri, F., Meng, Y., Schatzberg, A. F., 2017. Resting-state connectivity biomarkers define neurophysiological subtypes of depression. Nature medicine, 23(1), 28–38.

Elsabbagh M, Divan G, Koh YJ, Kim YS, Kauchali S, Marcin C, Montiel-Nava C, Patel V, Paula CS, Wang C, Yasamy MT, Fombonne E., 2012. Global prevalence of autism and other pervasive developmental disorders. Autism Res 5:160–179.

Filler AG., 2009. “The history, development, and impact of computed imaging in neurological diagnosis and neurosurgery: CT, MRI, DTI”. Nature Precedings.

Goldani, A. A., Downs, S. R., Widjaja, F., Lawton, B., & Hendren, R. L. (2014). Biomarkers in autism. Frontiers in psychiatry, 5, 100.

Grecucci, A., Siugzdaite, R., Job, R., 2017. Advanced Neuroimaging Methods for Studying Autism Disorder. Frontiers in neuroscience, 11, 533.

Ha, S., Sohn, I. J., Kim, N., Sim, H. J., Cheon, K. A., 2015. Characteristics of brains in autism spectrum disorder: structure, function and connectivity across the lifespan. Experimental neurobiology, 24(4), 273–284.

Huang, W., Goldsberry, L., Wymbs, N. F., Grafton, S. T., Bassett, D. S., Ribeiro, A., 2016. Graph frequency analysis of brain signals. J. Sel. Topics Signal Processing, 10, 1189–1203.

Huang, W., Bolton, T. A., Medaglia, J. D., Bassett, D. S., Ribeiro, A., Van De Ville, D., 2018. A graph signal processing perspective on functional brain imaging. Proceedings of the IEEE, 106(5), 868–885.

Hull, J. V., Jacokes, Z. J., Torgerson, C. M., Irimia, A., Van Horn, J. D., 2016. Resting-state functional connectivity in autism spectrum disorders: A review. Frontiers in Psychiatry, 7, 205.

Hull, J. V., Dokovna, L. B., Jacokes, Z. J., Torgerson, C. M., Irimia, A., Van Horn, J. D., 2017. Resting-state functional connectivity in autism spectrum disorders: A review. Frontiers in psychiatry, 7, 205.

Iidaka, T., 2015. Resting state functional magnetic resonance imaging and neural network classified autism and control. Cortex, 63, 55–67.

Itani, S., Thanou, D., 2019. Combining anatomical and functional networks for neuropathology identification: A case study on autism spectrum disorder. arXiv preprint arXiv:1904.11296.

Jou, R. J., Jackowski, A. P., Papademetris, X., Rajeevan, N., Staib, L. H., Volkmar, F. R., 2011. Diffusion tensor imaging in autism spectrum disorders: preliminary evidence of abnormal neural connectivity. Australian New Zealand Journal of Psychiatry, 45(2), 153–162.

Kolda, T. G., Bader, B. W., 2009. Tensor decompositions and applications. SIAM review, 51(3), 455–500.

Li, D., Karnath, H. O., Xu, X., 2017. Candidate biomarkers in children with autism spectrum disorder: a review of MRI studies. Neuroscience bulletin, 33(2), 219–237.

Mash, L. E., Reiter, M. A., Linke, A. C., Townsend, J., Müller, R. A., 2018. Multimodal approaches to functional connectivity in autism spectrum disorders: an integrative perspective. Developmental neurobiology, 78(5), 456–473.

Mash, L. E., Linke, A. C., Olson, L. A., Fishman, I., Liu, T. T., Müller, R. A., 2019. Transient states of network connectivity are atypical in autism: A dynamic functional connectivity study. Human brain mapping, 40(8), 2377–2389.

Medaglia, J. D., Huang, W., Karuza, E. A., Kelkar, A., Thompson-Schill, S. L., Ribeiro, A., Bassett, D. S., 2018. Functional alignment with anatomical networks is associated with cognitive flexibility. Nature Human Behaviour, 2, 156.

Mennigen, E., Miller, R. L., Rashid, B., Fryer, S. L., Loewy, R. L., Stuart, B. K., Calhoun, V. D., 2018. Reduced higher-dimensional resting state fMRI dynamism in clinical high-risk individuals for schizophrenia identified by meta-state analysis. Schizophrenia research, 201, 217–223.

Ménoret, M., Farrugia, N., Pasdeloup, B., Gripon, V., 2017. Evaluating graph signal processing for neuroimaging through classification and dimensionality reduction. In 2017 IEEE Global Conference on Signal and Information Processing (GlobalSIP) (pp. 618–622). IEEE.

Mukherjee, P., McKinstry, R. C., 2006. Diffusion tensor imaging and tractography of human brain development. Neuroimaging Clinics, 16(1), 19–43.

Nagae, L. M., Zarnow, D. M., Blaskey, L., Dell, J., Khan, S. Y., Qasmieh, S., Roberts, T. P. L., 2012. Elevated mean diffusivity in the left hemisphere superior longitudinal fasciculus in autism spectrum disorders increases with more profound language impairment. American Journal of Neuroradiology, 33(9), 1720–1725.

Nair, A., Treiber, J. M., Shukla, D. K., Shih, P., Müller, R. A., 2013. Impaired thalamocortical connectivity in autism spectrum disorder: a study of functional and anatomical connectivity. Brain, 136(6), 1942–1955.

Ortega, A., Frossard, P., Kovačević, J., Moura, J. M., Vandergheynst, P., 2018. Graph signal processing: Overview, challenges, and applications. Proceedings of the IEEE, 106(5), 808–828.

Power, J. D., Barnes, K. A., Snyder, A. Z., Schlaggar, B. L., Petersen, S. E., 2012. Spurious but systematic correlations in functional connectivity MRI networks arise from subject motion. Neuroimage, 59(3), 2142–2154.

Preprocessed Connectomes Project., 2014. ABIDE Preprocessed. http://preprocessed-connectomes-project.org/abide/. [Online; accessed 03-11-2018].

Preti, M. G., Bolton, T. A., Van De Ville, D., 2017. The dynamic functional connectome: State-of-the-art and perspectives. NeuroImage, 160, 41–54. https://doi.org/10.1016/j.neuroimage.2016.12.061.

Rashid, B., Blanken, L. M., Muetzel, R. L., Miller, R., Damaraju, E., Arbabshirani, M. R., Tiemeier, H., 2018. Connectivity dynamics in typical development and its relationship to autistic traits and autism spectrum disorder. Human brain mapping, 39(8), 3127–3142.

Shuman, D. I., Narang, S. K., Frossard, P., Ortega, A., Vandergheynst, P., 2013. The emerging field of signal processing on graphs: Extending high dimensional data analysis to networks and other irregular domains. IEEE Signal Processing Magazine, 30, 83–98.

Stankovic, L., Mandic, D., Dakovic, M., Brajovic, M., Scalzo, B., Constantinides, T., 2019. Graph signal srocessing – Part II: processing and analyzing signals on graphs. arXiv preprint arXiv:1909.10325v1.

Sundaram, S. K., Kumar, A., Makki, M. I., Behen, M. E., Chugani, H. T., Chugani, D. C., 2008. Diffusion tensor imaging of frontal lobe in autism spectrum disorder. Cerebral cortex, 18(11), 2659–2665.

Theiler, J., Eubank, S., Longtin, A., Galdrikian, B., Farmer, J. D., 1992. Testing for nonlinearity in time series: the method of surrogate data. Physica D: Nonlinear Phenomena, 58(1-4), 77–94.

Tzourio-Mazoyer, N., Landeau, B., Papathanassiou, D., Crivello, F., Etard, O., Delcroix, N., Joliot, M., 2002. Automated anatomical labeling of activations in SPM using a macroscopic anatomical parcellation of the MNI MRI single-subject brain. Neuroimage, 15(1), 273–289.

Uddin, L. Q., Supekar, K., Menon, V., 2013. Reconceptualizing functional brain connectivity in autism from a developmental perspective. Frontiers in.

Von Luxburg, U., 2007. A tutorial on spectral clustering. Statistics and computing, 17, 395–416.

Wang, J., Calhoun, V. D., Stephen, J. M., Wilson, T. W., Wang, Y. P., 2018. Integration of network topological features and graph Fourier transform for fMRI data analysis. In 2018 IEEE 15th International Symposium on Biomedical Imaging (ISBI 2018) (pp. 92–96). IEEE.

Woo, C. W., Wager, T. D., 2015. Neuroimaging-based biomarker discovery and validation. Pain, 156(8), 1379.

