## Supplementary file of paper for "Rest-fMRI Based Comparison Study between Autism Spectrum Disorder and Typically Control Using Graph Frequency Bands"

### S.1. Assigning AAL ROIs to seven well-known brain networks

Table S 1. Assigning 97 ROIs of AAL atlas to seven well-known brain networks including default mode network (DMN), cognitive control network (CCN), auditory network (AN), somatomotor network (SMN), visual network (VN), cerebral network (CN), subcortical network (SCN) . The L and R referred to left and right hemisphere of brain.

| | | | | |
|---|---|---|---|---|
| **DMN** | Frontal_Mid_R | Temporal_Inf_L | Calcarine_L | Cerebelum_4_5_L |
| Frontal_Sup_Medial_L | Frontal_Mid_Orb_L | Temporal_Inf_R | Calcarine_R | Cerebelum_4_5_R |
| Frontal_Sup_Medial_R | Frontal_Mid_Orb_R | **AN** | Cuneus_L | Cerebelum_6_L |
| Frontal_Med_Orb_L | Frontal_Inf_Oper_L | Heschl_L | Cuneus_R | Cerebelum_6_R |
| Frontal_Med_Orb_R | Frontal_Inf_Oper_R | Heschl_R | Lingual_L | Cerebelum_7b_L |
| Cingulum_Ant_L | Frontal_Inf_Tri_L | Temporal_Sup_L | Lingual_R | Cerebelum_8_L |
| Cingulum_Ant_R | Frontal_Inf_Tri_R | Temporal_Sup_R | Occipital_Sup_L | Cerebelum_8_R |
| Cingulum_Mid_L | Frontal_Inf_Orb_L | **SMN** | Occipital_Sup_R | Cerebelum_9_L |
| Cingulum_Mid_R | Frontal_Inf_Orb_R | Precentral_L | Occipital_Mid_L | Cerebelum_9_R |
| Cingulum_Post_L | Insula_L | Precentral_R | Occipital_Mid_R | Cerebelum_10_L |
| Cingulum_Post_R | Insula_R | Supp_Motor_Area_L | Occipital_Inf_L | Cerebelum_10_R |
| Angular_L | Hippocampus_L | Supp_Motor_Area_R | Occipital_Inf_R | **SCN** |
| Angular_R | Hippocampus_R | Olfactory_L | Fusiform_L | Caudate_L |
| Precuneus_L | ParaHippocampal_L | Olfactory_R | Fusiform_R | Caudate_R |
| Precuneus_R | ParaHippocampal_R | Postcentral_L | **CN** | Putamen_L |
| **CCN** | Parietal_Sup_L | Postcentral_R | Cerebelum_Crus1_L | Putamen_R |
| Frontal_Sup_L | Parietal_Sup_R | SupraMarginal_L | Cerebelum_Crus1_R | Pallidum_L |
| Frontal_Sup_R | Parietal_Inf_L | SupraMarginal_R | Cerebelum_Crus2_L | Pallidum_R |
| Frontal_Sup_Orb_L | Parietal_Inf_R | Paracentral_Lobule_L | Cerebelum_Crus2_R | Thalamus_L |
| Frontal_Sup_Orb_R | Temporal_Mid_L | Paracentral_Lobule_R | Cerebelum_3_L | Thalamus_R |
| Frontal_Mid_L | Temporal_Mid_R | **VN** | Cerebelum_3_R | |

### S.2. Some of the QCC analyses

For the first analysis, the probability of co-occurring of two levels existing in first and eighth bands are investigated. Let us express more clearly this analysis by an example. For first band, the time points at which level 1 is occurred are determined (denoted by *TT1*). For each level of the eighth band, the probability of co-occurring is computed by dividing the number of its occurring at *TT1* to the number of *TT1* time points. This is implemented for all levels of first band and also the same approach is performed for the eighth frequency band when computing its levels co-occurring with levels of first band. The results are shown in Fig.S1.

For the second analysis, the *effect* of increasing/decreasing CC level of one band on the other band increasing/decreasing is investigated. To do this for a given subject, for first/eighth band, the difference between consecutive CC level values is computed in order to obtain >0, 0, or <0 for increasing, unchanging, or decreasing the CC level, respectively. Then, for first/eighth band, the time points at which increasing/decreasing is occurred are determined (denoted by *TT_inc/TT_dec*). For eighth/first band, the time points [*TT_inc*, *TT_inc+1*] are investigated for finding the number of increasing (denoted by *Num_inc*) and the number of decreasing (denoted by *Num_dec*). The *Num_inc* and *Num_dec* are divided to the number of *TT_inc* time points in order to have normalized *effect* value. For obtaining the *Num_inc* and *Num_dec* one time point after each time point of *TT_inc* is also sought because it may be possible that level changing in the eighth/first band is started with one TR delay. The same process is also performed for [*TT_dec*, *TT_dec+1*]. The results are shown in Fig.S 2.

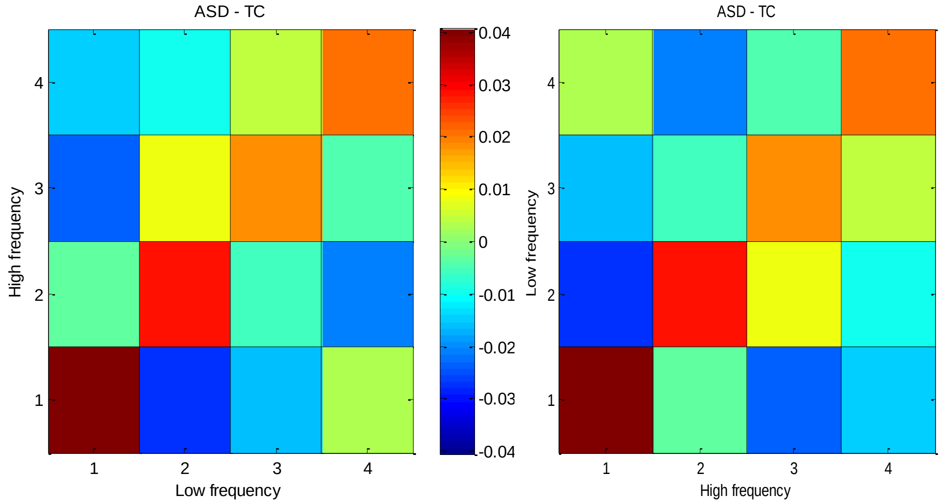


Fig.S1. The mean probability of co-occurring for ASD group subtracted from that of TC group. The left/right side is for the probability of co-occurring between each CC level at high/low frequency band and the given CC level at low/high frequency band.

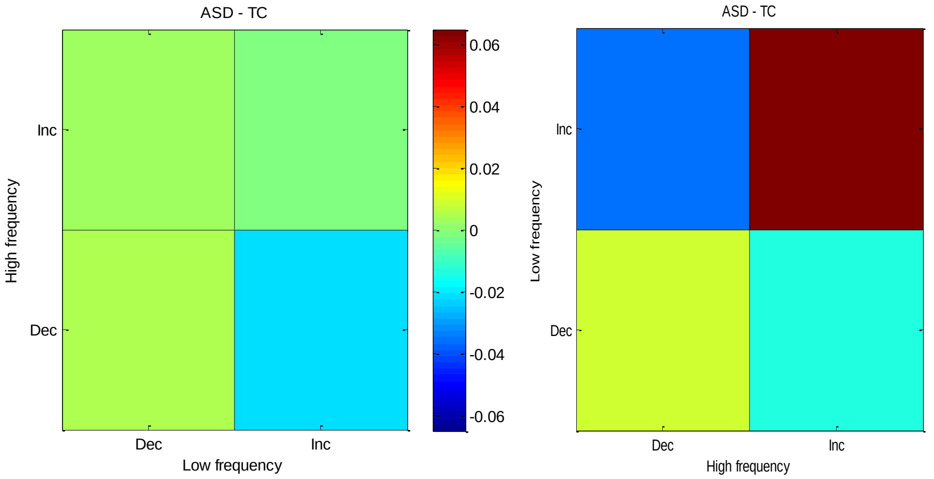


Fig.S 2. The effect of increasing/decreasing CC levels at the low frequency band on the increasing and decreasing CC levels at the high frequency band (left side); similar analysis for CC level changes at low frequency band with respect to CC level changes at high frequency band (right side).

### S.3. Some results of within and between networks FC

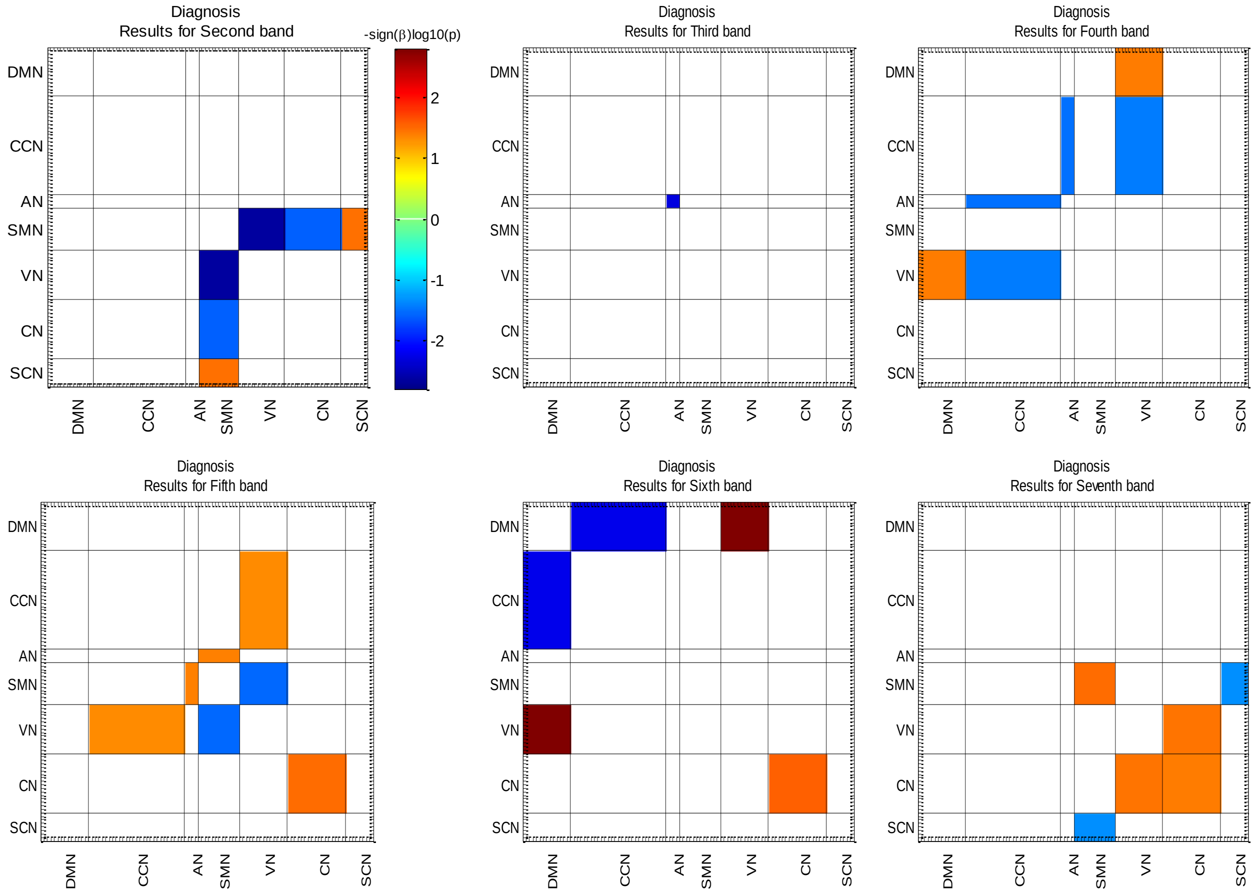


Fig.S 3. Results of FC analysis for diagnosis variable at second, third, fourth, fifth, sixth, and seventh frequency bands. All colored connectivity are significant ($p < 0.05$).

### S.4. Elbow criterion for k-means clustering of LVFS analysis

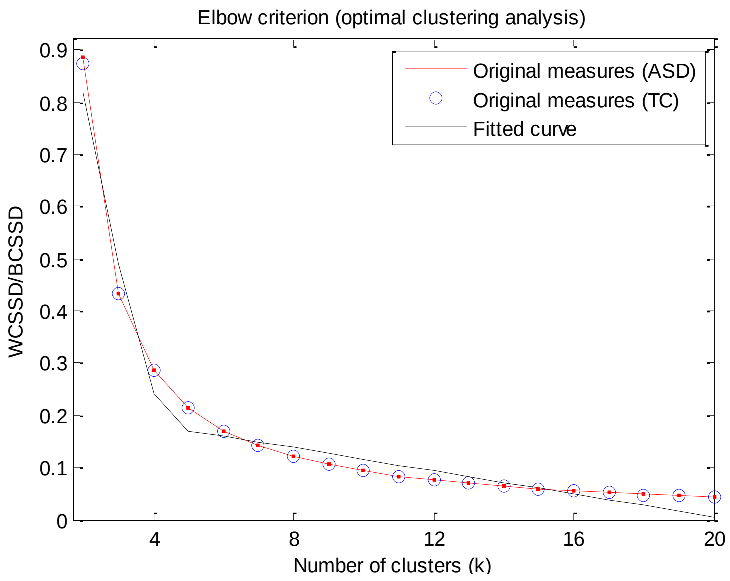


Fig.S 4. The Elbow criterion for selecting the optimum number of clusters of LVFS analysis. The red line and blue circles show the original measures of ratio WCSD/BCSSD for ASD and TC with respect to the number of clusters (k), and the black line represents the best fitted elbow-shaped curve to the ASD and TC original measures. The optimum cluster number is 5.

### S.5. Cross frequency FC matrices

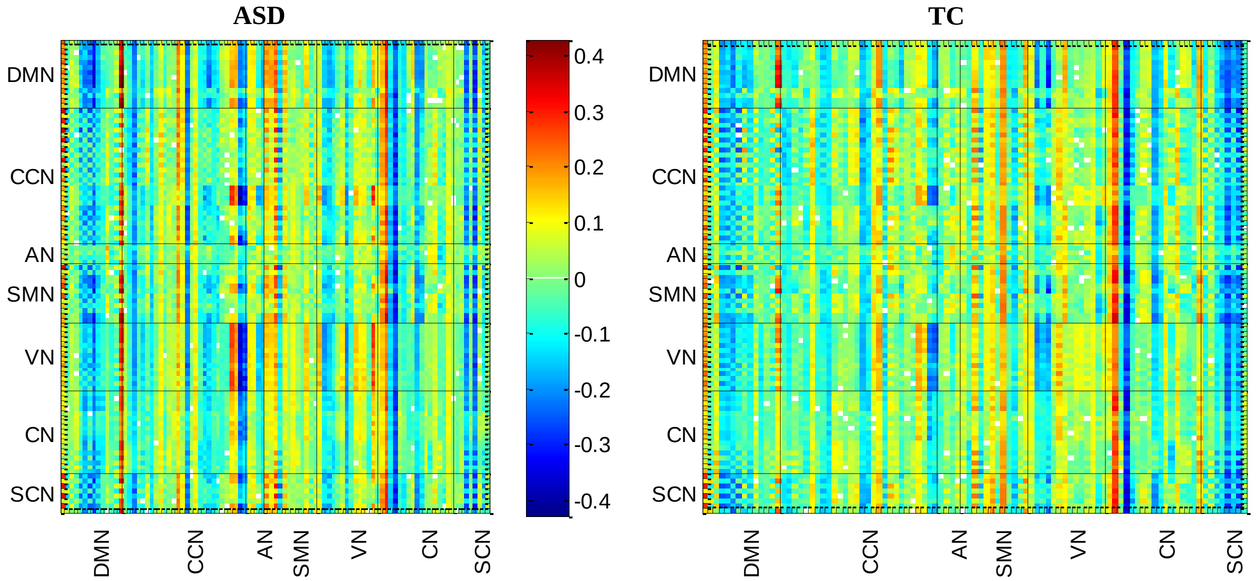


Fig.S 5. The z-scored cross frequency FC of ASD and TC groups are illustrated. The matrix of each group is obtained by averaging across that group subjects.

### S.6. Cross frequency FC values of some important diagnostic ROIs

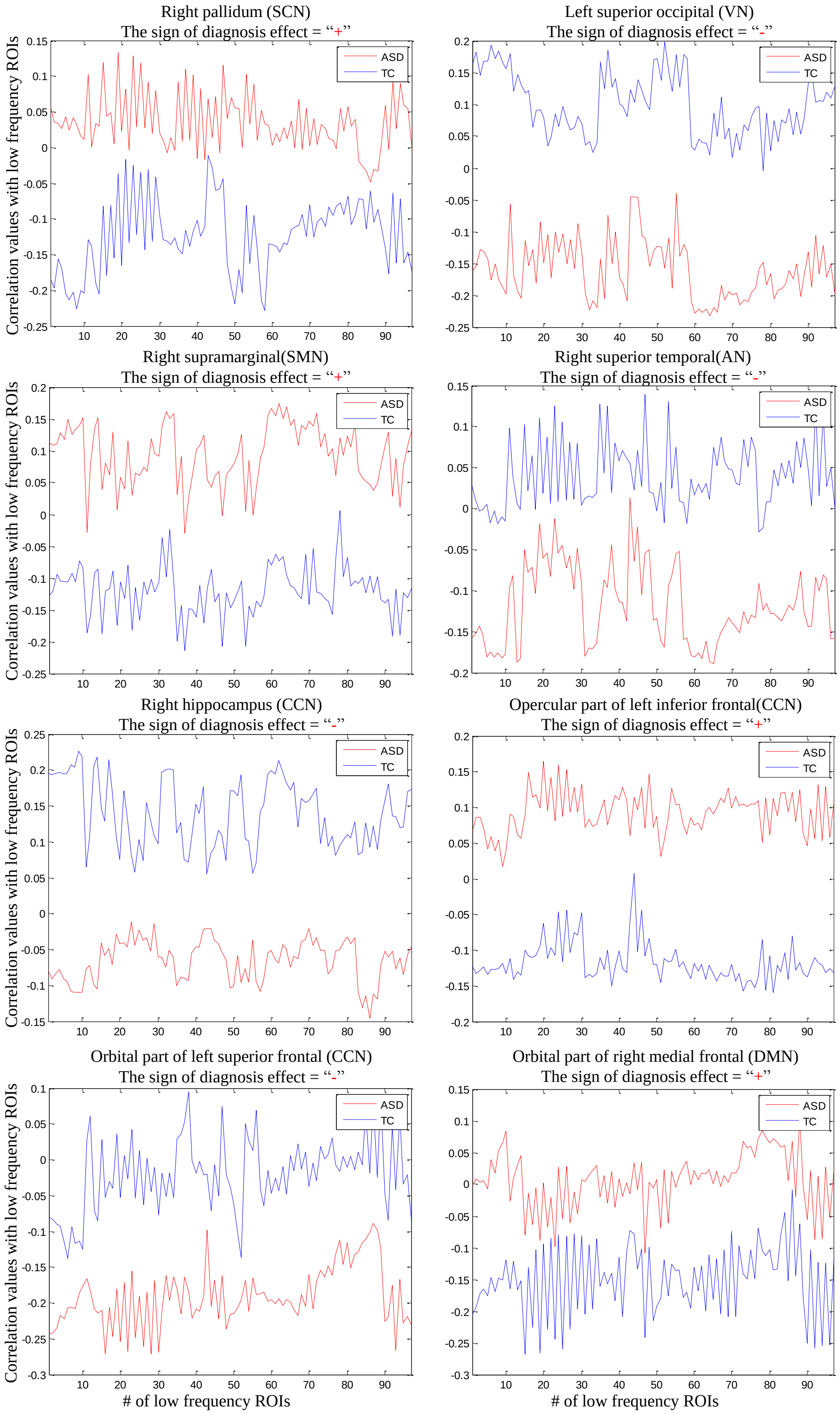


Fig.S 6. Each plot shows the FC values that high frequency signal of one ROI has with low frequency signals of 97 understudy ROIs. The first row of title indicates the name of high frequency ROI and its corresponding network and the second column presents the dominant diagnosis effects sign observed for that high frequency ROI. All of these plots are for ROI levels analysis of ASD and TC groups and the mentioned FC values are obtained by averaging across group subjects.

### S.7. Tensor factorization of ASD and TC groups

#### *S.7.1. Method and implementation*

In this section of supplementary file, the detailed information of performing tensor factorization (TF) and its results are reported. A 3D tensor for ASD and TC data can be separately employed where the first, second and third dimensions stand for FC vector, subjects, and graph frequency bands, respectively. This analysis is performed at ROI and network levels. At ROI level, the FC matrices of subjects are obtained as explained for third scenario (subsection 2.4.3) of paper and then the age and sex effects are regressed out from each ROI-ROI FC values. Finally, the upper triangular entries of FC matrices (due to symmetric property of FC matrices) are used as first dimension. At network level, the network FC matrices are computed as explained for third scenario (subsection 2.4.3) and then the age and sex effects are removed from within and between network FC values. Finally, the upper triangular and diagonal entries of network FC matrices are selected as first dimension of 3D tensor.

A TF technique aims at introducing one factor for each dimension of tensor so that that factor represents the most variance and important information of its corresponding dimension. Two well-known techniques for TF of a tensor are used: Tucker decomposition, canonical polyadic (CP) decomposition (Kolda and Bader, 2009). The tucker decomposition outputs are factor matrices which are orthogonal and a core matrix. The entries of core matrix represent the weight of interactions existing between factors of different dimensions. In Tucker decomposition, the number of columns (components) of each factor can be different from those of other factors. The CP output is factor matrices and CP components can be linearly dependent. All factors of CP decomposition have the same number of components $R$. In fact, the CP decomposition factorizes a tensor into a sum of component rank-one tensors (Kolda and Bader, 2009). It has been shown that the Tucker decomposition is more suitable for classification studies and CP decomposition is more suitable for studies needing interpretation (Kolda and Bader, 2009, Mokhtari et al., 2018). Hence, in this analysis, the CP decomposition is considered.

One of the most important step of TF implementation is determining the proper number of components of factors $R$. In this study, the core consistency technique is employed for determination of $R$ (Bro and Kiers et al., 2003). For TF analyses, the N-way toolbox is used (Andersson and Bro, 2000). After TF of ASD and TC data, the correlation of ASD graph frequency bands components with those of TC are computed and the best matched components of two groups are found. Based on this matching result, the components of other factors of ASD and TC are ordered.

#### *S.7.2. Results*

The results of core consistency are shown in Fig.S 7. It is obvious that proper $R$ values of ASD and TC at network level and TC at ROI level are 2, 2, and 3, respectively. The ASD at ROI level obtains core consistency values of [100, 99, 74.98, 66, 35.6] for number of components [1, 2, 3, 4, 5], respectively. Sharp drop occurs at $R$=3 but even for $R$=3 and 4 the core consistency values are very high and so the four-component model may be the best choice. This behavior of core consistency values maybe show that the ASD data is difficult to model adequately in comparison to the TC one.

The ASD and TC factors at network and ROI levels are shown in Fig.S 8 and Fig.S 9, respectively. At network level, only positive relations between networks exist in first component of ASD and TC. In this component, the DMN connections with all of the rest networks is more for ASD than TC. For TC, the DMN connection with VN is much stronger than SCN whereas for ASD the DMN connection with SCN is stronger than The VN. VN-VN connection for TC is very strong in component1. The third factor results show that first component is happened only in low frequency band (GFB1) for both ASD and TC. In second component, both positive and negative connections exist for ASD and TC. The most distinct difference between ASD and TC is the connection between AN and SCN which is strongly positive for ASD and weakly negative for TC. By considering the third factor, it can be said that the second component presence of ASD in GFB1 is much stronger than TC. In middle bands, two groups show somewhat similar behavior. However, in high frequency bands (GFB7 and GFB8), two groups show contradictory behavior. The ASD increases its positive presence in these bands whereas the TC increases its negative presence in these bands. In brief, in second component, ASD and TC follow/contradictory follow each other in low/high frequency bands. As a result, it can be said that AN-SCN is negative and positive for ASD and TC in low frequency bands and negative for both ASD and TC in high frequency bands.

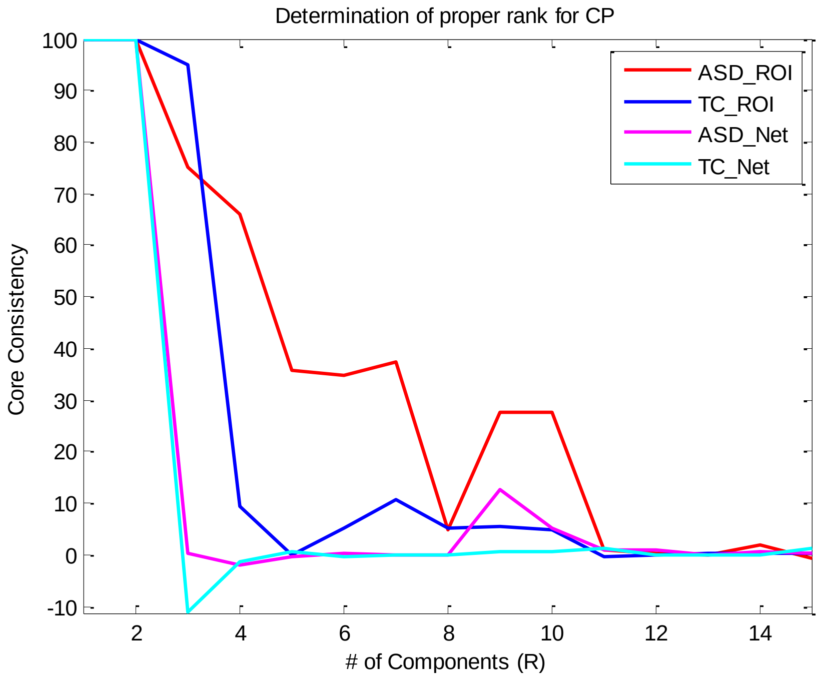


Fig.S 7. Core consistency analyses for ASD and TC at ROI and network (Net) levels. The best $R$ value for both ASD and TC at network value is 2. The $R$ for TC at ROI level is 3. The core consistency values of ASD at ROI level show sharp drop after $R>2$, but even after this drop its values are still high (0.75 for $R=3$ and 0.66 for $R=4$). Thus, the best $R$ value for ASD at ROI level is 4.

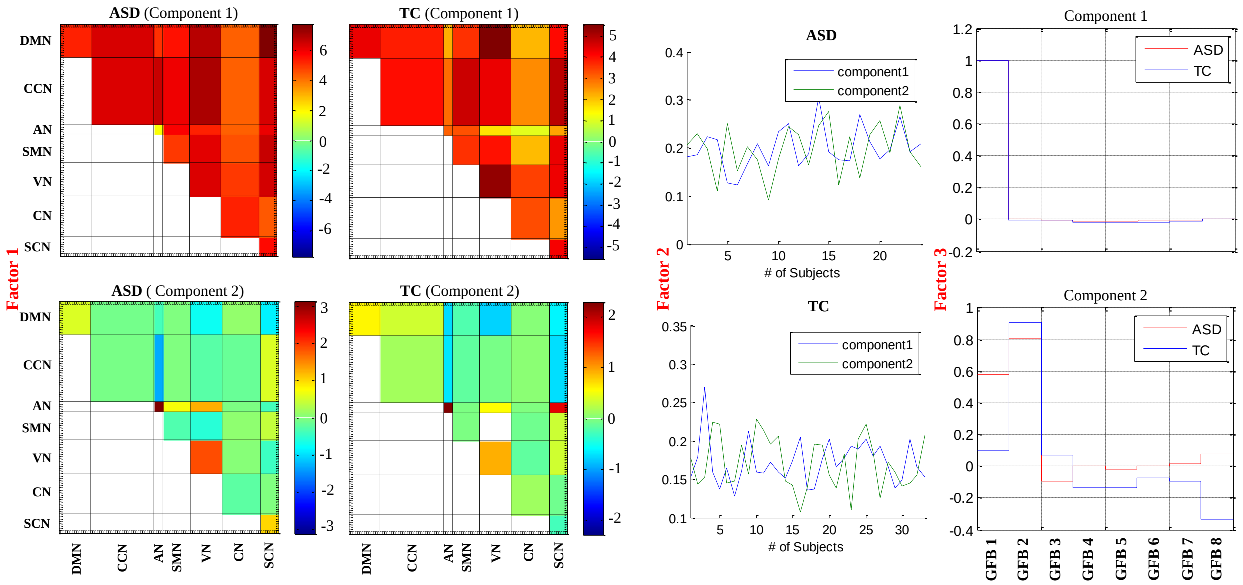


Fig.S 8. Tensor factorization results of network level analyses. The components of three factors which are related to FC matrices, subjects, and graph frequency bands (GFB)s of ASD and TC are shown.

At ROI level, positive ROI-ROI connections are dominant for both ASD and TC in first component. The positive ROI-ROI connections are much more and much stronger and much clustered form for ASD in comparison to TC. The second, third and fourth components of first factor of ASD are similar to each other and are dominated by very week or close to zeros connections. However, some strong positive and negative ROI-ROI connections are seen in CN-CN. In second/third component, some strong negative/strong positive ROI-ROI connections exist in VN-VN and CCN-CCN. For TC, the second and third components of first factor are very similar to each other (correlation value of 0.83) but distinct difference between these two components can be seen for ROI-ROI connections of CN-CN where strong

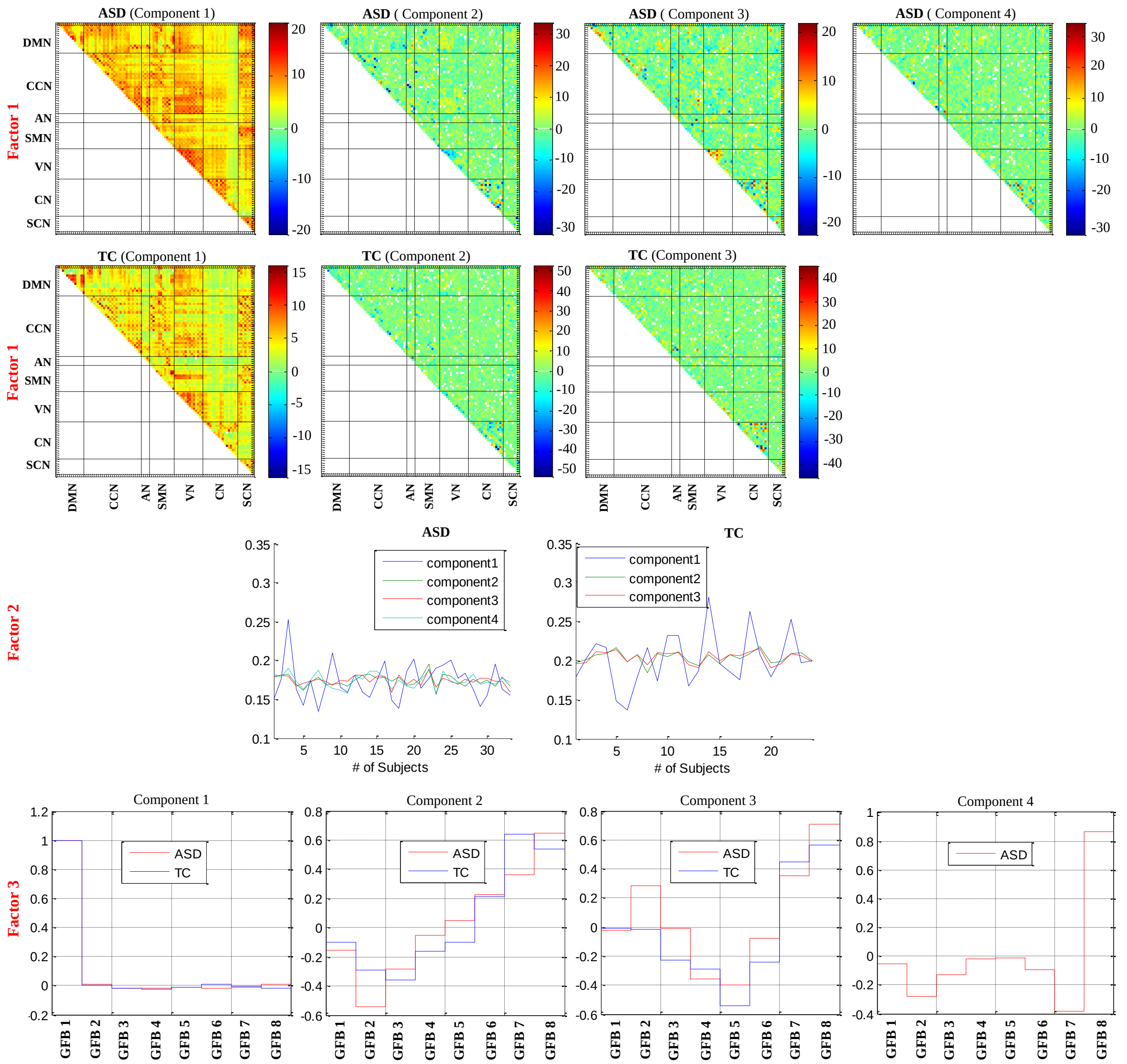


Fig.S 9. Tensor factorization results of ROI level analyses. The components of three factors which are related to FC matrices, subjects, and graph frequency bands (GFB)s of ASD and TC are shown.

negative/strong positive connections are dominant for second/third component.

For the third factor, as with network level, the first component exists in GFB1 for both ASD and TC. The second and third components of ASD and TC strongly follow each other (except in the GFP8 of second component which is high frequency band). The second component is decreased in GFB2 and is increased in GFB3-GFB8 whereas the third component is decreased GFB3 and GFB5 and is increased in GFB6-GFB8. The presence of component 4 is strong in GFB8 and in middle GFBs and GFB1 is very low.

The components of second factor have positive values for both ASD and TC at ROI and network levels. The first component of this factor has dominant amplitude in comparison to other components at ROI level. However, at network level this dominancy cannot be seen. It is maybe that the second component of network level show itself in two (for TC) and three (for ASD) different components at ROI level and so it’s corresponding amplitude is divided between two (for TC) and three (for ASD) components of ROI level.

### References


Andersson, C. A., and Bro, R. 2000. The N-way toolbox for MATLAB. Chemometrics and intelligent laboratory systems, 52(1), 1-4.

Bro, R., and Kiers, H. A. 2003. A new efficient method for determining the number of components in PARAFAC models. Journal of Chemometrics: A Journal of the Chemometrics Society, 17(5), 274-286.

Kolda, T. G., Bader, B. W., 2009. Tensor decompositions and applications. SIAM review, 51(3), 455-500.

Mokhtari, F., Laurienti, P. J., Rejeski, W. J., Ballard, G. 2018. Dynamic fMRI connectivity tensor decomposition: a new approach to analyze and interpret dynamic brain connectivity. Brain Connect, 9, 95-112.
